# A robust human airway organoid platform enables scalable expansion and trajectory mapping of pulmonary neuroendocrine cells

**DOI:** 10.64898/2026.01.10.698835

**Authors:** Noah Candeli, Lisanne den Hartigh, Nicholas Hou, Andrés Marco, José Antonio Sánchez-Villicaña, Andrea García-González, Shashank Gandhi, Francesca Sgualdino, Alyssa J. Miller, Jason Spence, Susana Chuva de Sousa Lopes, José L. McFaline-Figueroa, Hans Clevers, Talya L. Dayton

## Abstract

Pulmonary neuroendocrine cells (PNECs) are rare chemosensory epithelial cells, facultative stem cells, and a cell-of-origin for neuroendocrine lung cancers, yet the mechanisms governing their differentiation and heterogeneity are poorly understood. Here we establish NEr-fAOs, a human fetal airway organoid platform that robustly enriches PNECs, and identify a cooperative requirement for dual GSK3 and NOTCH inhibition to drive directed PNEC differentiation. This strategy yields stable cultures with up to 60-fold expansion of PNECs whose transcriptomes closely match fetal and adult PNECs. In addition to PNEC-enrichment, NEr-fAOs retain diverse airway epithelial cell types, preserving epithelial complexity. Time-resolved single-cell transcriptomics maps PNEC trajectories in NEr-fAOs, resolving precursor and mature states. Comparative analyses further reveal a distal airway bias in NEr-fAOs and enrichment for lower-airway progenitors. NEr-fAOs thus provide a scalable, tractable platform to dissect human PNEC biology and distal airway progenitor hierarchies relevant to lung development, cancer, and disease.

## INTRODUCTION

Pulmonary neuroendocrine cells (PNECs) are a rare, evolutionarily conserved population of airway epithelial cells comprising approximately 0.5% of the lung epithelium^1^. Despite their low abundance, PNECs are increasingly recognized as critical regulators of lung homeostasis, contributing to chemosensing^2,3^, mechanosensation^4^, epithelial regeneration^5,6^, and immune regulation^7^. Aberrant PNEC function and altered abundance have been reported in several chronic pulmonary diseases, including cystic fibrosis, asthma, and chronic obstructive pulmonary disease^8,9^. Moreover, PNECs are considered a primary cell of origin for small-cell lung cancer (SCLC), which accounts for 25% of aggressive lung cancers^10,11^, and serve as the presumptive cells of origin for other subtypes of neuroendocrine lung cancers, such as the less common large-cell neuroendocrine carcinoma (LCNEC) and carcinoids.

PNECs have an intrinsic epithelial origin^12^ and are the first differentiated cell type to appear in the developing lung^13^. *In vitro* studies show that PNECs originate from Lower Airway Progenitor (LAP) cells, a population enriched in the lower small cartilaginous and noncartilaginous airways (NCA) of the fetal lung, arising during branch morphogenesis^14^. Mouse studies provide evidence that PNECs in the trachea can originate from basal cells^15^.

Efforts to study PNECs have been hampered by their extreme rarity, lack of robust isolation methods, and the difficulty of sustaining human airway epithelial cells in long-term culture. Nonetheless, multiple *in vitro* models have been developed to address these challenges, generating PNECs under diverse culture conditions using both pluripotent stem cells (PSCs) and tissue-resident stem cells (TSCs)^16–21^. Despite their utility, PSC-derived systems are constrained by lengthy differentiation protocols and limited potential for long-term expansion. In contrast, models derived from TSCs provide a platform for the long-term expansion and differentiation of the human airway epithelium *in vitro*. ^22^ and ^23^ were the first to generate TSC-derived airway organoids respectively from fetal and adult human lungs. However there was no reported presence of PNECs in these models. ^20^ were the first to report PNEC generation from tissue-stem cell derived 3D bud tip airway organoids differentiated through SMAD inhibition. ^18^ reported generation of PNECs from human tracheal epithelium differentiated in 2D ALI culture. ^21^ generated 2D ALI and 3D organoid cultures from primary human distal lung tissue grown in Standard Airway Organoid medium ^23^ or commercial media from STEMCELL Technologies which included rare epithelial cell types including PNEC, tuft cells and ionocytes. Although these organoid systems allow for the long-term expansion of the airway epithelial cells that can differentiate into PNECs and other relevant cell types (basal, club, multiciliated, goblet), PNEC abundance in these systems remains low, limiting their utility for mechanistic studies or modeling PNEC-driven diseases. Moreover the precise *in vitro* conditions responsible for driving the specification of lung progenitors into PNEC precursors and their subsequent maturation into differentiated PNECs remains unclear.

Here, we report a scalable *in vitro* model for the expansion and direct differentiation of human fetal airway organoids (fAOs) enriched in PNECs, NEr-fAOs. By optimizing a neuroendocrine (NE) expansion medium, we established long-term cultures containing PNECs and their progenitors. Subsequent differentiation protocols yielded organoids with high PNEC abundance. Single-cell RNA sequencing confirmed a high degree of transcriptomic similarity between *in vitro*-derived epithelial cells and their *in vivo* counterparts while also revealing previously uncharacterized heterogeneity and specification mechanisms guiding PNEC development. This platform enables detailed molecular interrogation of human PNEC biology and provides a foundation for modeling neuroendocrine-related lung diseases.

## RESULTS

### Generation of neuroendocrine enriched NEr-fAOs

Our previously described adult airway organoid (AO) cultures enable long-term expansion of organoids containing multiple airway epithelial cell types, but consistently lack pulmonary neuroendocrine cells (PNECs) ^23^. Because PNECs make up only ∼0.5% of the lung epithelium^1^, their absence could reflect either low starting abundance or selective loss during culture. To generate organoids containing all airway epithelial cells, including PNECs, we established fetal AOs (fAOs) from healthy human lungs at gestational weeks (GW) 15-19. fAOs were generated by enzymatic digestion of whole lung lobes and cultured in our published AO media (“standard” media). In contrast to earlier fAO protocols, we used the entire lung rather than only the distal tip, thereby avoiding bias towards a particular progenitor cell population ^22,24^.

RT-qPCR analysis of the PNEC-lineage transcription marker, *ASCL1,* revealed expression at passage 2 (P2) of our fAOs but a marked decline to nearly undetectable levels by P6-7 (Fig. S1a), suggesting that PNECs are lost during *in vitro* expansion in standard media, likely due to missing niche cues. To identify conditions that would sustain PNECs in fAOs, starting at isolation, we grew fAOs in either standard media or in standard media supplemented with growth factors previously linked to neuroendocrine growth (Fig. 1a). We tested five supplements: (1) midkine, (2) EGF, (3) FGF2, (4) the GSK3 inhibitor, CHIR99021 (CHIR), or (5) CHIR and FGF2 (CF2) ^25–27^ (Fig. S1b). Only fAOs grown in CHIR- or CF2-supplemented media showed sustained expression of *ASCL1,* reaching levels (normalized to whole lung tissue) of 113 to 348, compared to 0.5 to 37 in unsupplemented media.

**Figure 1.**
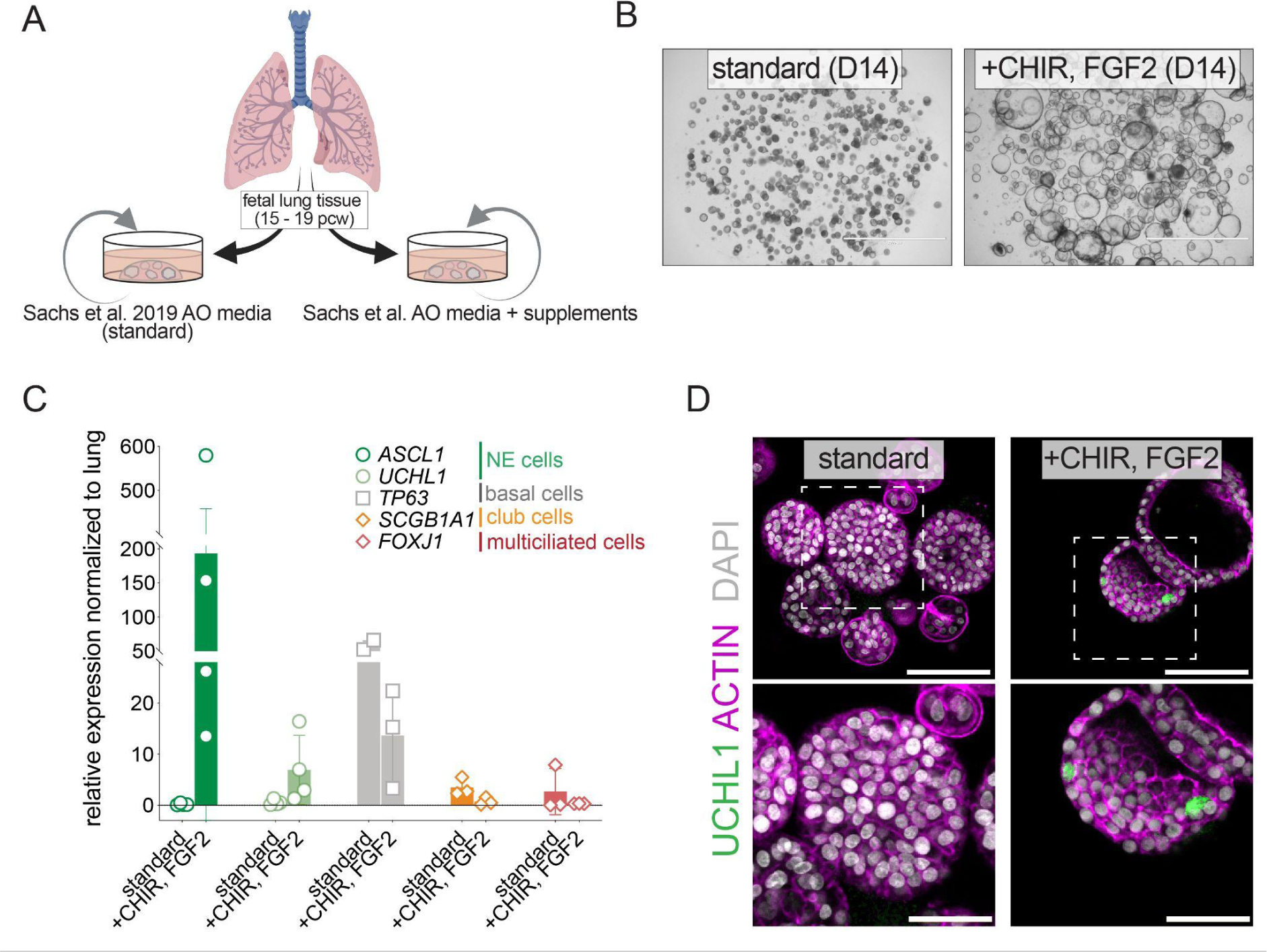
Optimised NE Expansion medium allows PNEC generation in fAOs. (A) Schematic of experimental design for testing growth factor sustaining PNECs in fAOs. (B) Brightfield images of fAOs organoids cultured for 14 days with indicated factors. Scale bar, 2000 μM (C) qPCR analysis of PNEC (ASCL1,UCHL1), basal (TP63), Club (SCGB1A1), and multiciliated (FOXJ1) markers in fAOs at P6 cultured with indicated factors for 14 days. Data are represented as mean ± SD and correspond to 4 different lines. (D) IF image of fAOs in standard or CF2. ACTIN = membrane stain; UCHL1 = PNEC marker. Scale bars 100 μM for top panels and 50 μM for bottom panels.

Because CF2 supported slightly faster growth than CHIR alone, we adopted it for subsequent experiments. fAOs in CF2 displayed distinct morphology and took longer to reach P5 than donor-matched lines in standard media *(average 92 vs. 61.4 days, respectively)* (Fig. S1c). While fAOs in standard media tended to be smaller, did not always form lumens, and had thick walls, fAOs in CF2 formed large, cystic organoids with thin walls (Fig. 1b).

To determine whether CF2 fAOs maintain PNECs alongside other airway epithelial cell types, we measured the expression of markers for PNECs (*ASCL1*, *UCHL1*), basal cells (*TP63*), club cells (*SCGB1A1*), and multiciliated cells (*FOXJ1*) in four donor-derived lines grown in either standard or CF2 (Fig. 1c). Whereas PNEC marker expression was undetectable in fAOs in standard media at P6, it was consistently high in fAOs in CF2 (p=0.03 by Mann Whitney U test for *ASCL1*). Compared to fAOs in standard media, fAOs in CF2 also showed a trend towards lower expression of the basal cell marker, *TP63*, pointing to potential differences in basal cell dynamics. These patterns were also observed at the later passage of P9 (Fig. S1d). Immunofluorescence (IF) staining using an antibody for the PNEC marker, UCHL1, confirmed the presence of PNECs in fAOs derived from multiple unrelated donors in CF2 media but not when grown in standard media (Fig. 1d and S1e). Based on these results, we termed CF2 media, Neuroendocrine (NE) Expansion media and the resulting PNEC-containing fAOs NE-enriched fAOs (NEr-fAOs).

### Dual inhibition of GSK3 and NOTCH signaling drives directed PNEC differentiation

NE Expansion media (CF2) supported long-term proliferation and self-renewal of NEr-fAOs. However, qRT-PCR showed that PNEC marker expression levels varied across passages, and IF for PNEC markers showed that these cells remained rare (Figure 1). We therefore developed a differentiation protocol to further enrich and mature PNECs in NEr-fAOs.

Notch signaling is a well-established repressor of the PNEC fate in the developing mouse lung, and Notch signaling inhibitors have been used in other *in vitro* systems to induce PNECs ^9,16,17,19,24^. This, together with the observation that only expansion media containing CHIR enabled the generation of NEr-fAOs, led us to test Notch inhibition and GSK3 inhibition (Fig. 1). We allowed NEr-fAOs to grow in NE Expansion media for 20 to 30 days, and then switched the media to one of three differentiation conditions: a growth-factor low base media supplemented with either the Notch inhibitor, DAPT, CHIR, or both (Fig. 2a). The time at which differentiation media was added corresponds to d0 of differentiation, day 20-30 of expansion.

**Figure 2.**
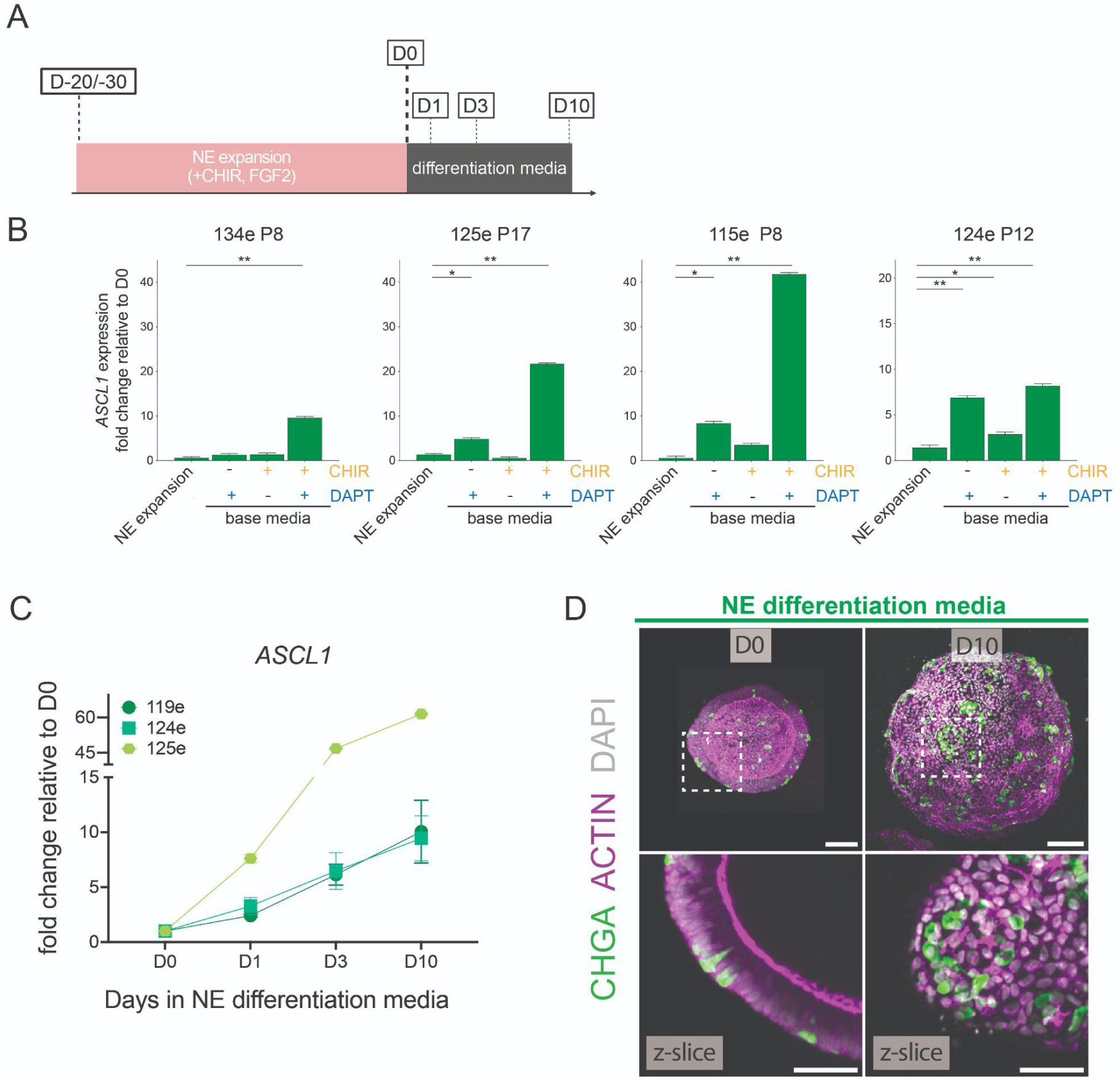
Dual inhibition of GSK3 and NOTCH signaling promotes PNEC differentiation. (A) Schematic of Neuroendocrine differentiation protocol in fAOs. (B) qPCR analysis of PNEC (*ASCL1*) marker in fAOs cultured with indicated factors for 20-30 days. Data are represented as mean ± SD. *P*-values using unpaired *t*-test are shown. (*) *P* < 0.05; (**) *P* < 0.01; (***) *P* < 0.001. (C) qPCR analysis of PNEC (ASCL1) marker in fAOs cultured with indicated factors for 20-30 days. Data are represented as mean ± SD. (D) IF image of fAOs in standard or CF2. ACTIN = membrane stain; UCHL1 = PNEC marker. Scale bars 100 μM for top panels and 50 μM for bottom panels.

In 3 out of 4 lines, DAPT alone induced *ASCL1* expression after 10 days of treatment *(5-8 fold vs. day 0 of differentiation)* (Fig. 2b). However, in all lines tested, dual DAPT+CHIR led to the strongest induction of *ASCL1* expression in NEr-fAOs (*8-40-fold induction vs. day 0*), pointing to potential synergy between dual inhibition of GSK3 and Notch signalling in promoting the PNEC fate. The same pattern was observed for an additional 2 PNEC markers, *UCHL1* and *NEUROD1* (Fig. S2a,b). Decreased expression of the Notch target gene, *HES1*, was seen in all DAPT conditions, confirming effective Notch blockade (Fig. S2c).

To investigate PNEC differentiation dynamics, we performed a qRT-PCR time course for 10 days after switching to base media + DAPT and CHIR, hereafter referred to as NE Differentiation media (Fig. 2c). This analysis revealed rapid *ASCL1* upregulation within 1 day in NE differentiation media, with continued increase over 10 days and reaching up to 60-fold increase over day 0. IF staining for the mature PNEC marker, CHGA, confirmed the mRNA results, revealing more PNECs in NEr-fAOs cultured in NE Differentiation media for 10 days than at day 0 (Fig. 2d and Fig. S2d). CHGA+ PNECs in differentiated NEr-fAOs appeared as either isolated cells or small clusters reminiscent of the neuroendocrine bodies observed at airway branchpoint in endogenous lung tissue ^13,28,29^. Altogether, these experiments demonstrate that dual GSK3 and Notch inhibition enables directed and scalable PNEC differentiation.

### Transcriptomic characterization of NEr-fAOs

To interrogate the cell composition of fAOs grown in the different media conditions, and evaluate their transcriptomic profile, we performed single-cell RNA sequencing using SORT-seq^30^. Samples were sorted from cultures maintained for 14 - 21 days in either standard media or NE Expansion media (NEr-fAOs). Samples were also harvested from NEr-fAOs in NE Differentiation, at various time points during differentiation (Day 1, 3, and 10) (Fig. 3a). In total, after quality control filtering, we sequenced and analyzed 5,540 fAO and NEr-fAO-derived cells. Leiden clustering analysis of the full dataset identified 13 clusters, with 3 clusters (clusters 1, 7, and 11) unique to NEr-fAOs (Fig. S3a,b,c, Table S1). To assign cell identities we performed automated cell type annotation using CellTypist^31^. We trained a custom CellTypist model using previously published scRNA-Seq datasets from human fetal^24^ and adult lung epithelium^32^ as reference. This enabled the identification of Basal, Secretory, Bud Tip Adjacent, RAS, Goblet, Multiciliated, and PNEC populations (Fig. S3d,e). To validate these annotations, we integrated our dataset with the reference atlases and observed that fAO cells closely aligned with those from lung epithelial datasets (Fig. 3b,c). Further validation using cell-type signature scores based on known fetal and adult lung markers confirmed the identities of these populations (Fig. S3f,g). We found that cells annotated as respiratory airway secretory (RAS) cells had high expression of both the RAS signature and the signature for the fetal airway stem cell population, lower airway progenitor (LAP) cells. We therefore named these cells RAS/LAP-like. Cells that had been annotated as alveolar type 1 (AT1) and bud tip adjacent had high expression of signatures from multiple progenitor cells including LAP, RAS, bud tip adjacent, and bud tip progenitor, as well as some more differentiated cell types. We therefore labelled these cells jointly as transitional distal progenitor-like cells (Fig 3d). Analysis of cell type contribution showed fAOs in all culture conditions retain airway cells such as basal, club, and multiciliated cells. However, PNECs and goblet cells were most abundant in NEr-fAOs in NE Differentiation media. Notably, the proportion of PNECs increased from ∼5% under NE expansion conditions to ∼12% after 10 days of NE differentiation (Fig. 3e and S3h). Interestingly, compared to fAOs in standard media, NEr-fAOs in NE expansion and NE differentiation are enriched for RAS/LAP-like cells. LAP cells are a fetal progenitor population that has been shown to give rise to PNECs during human fetal lung development^14^.

**Figure 3.**
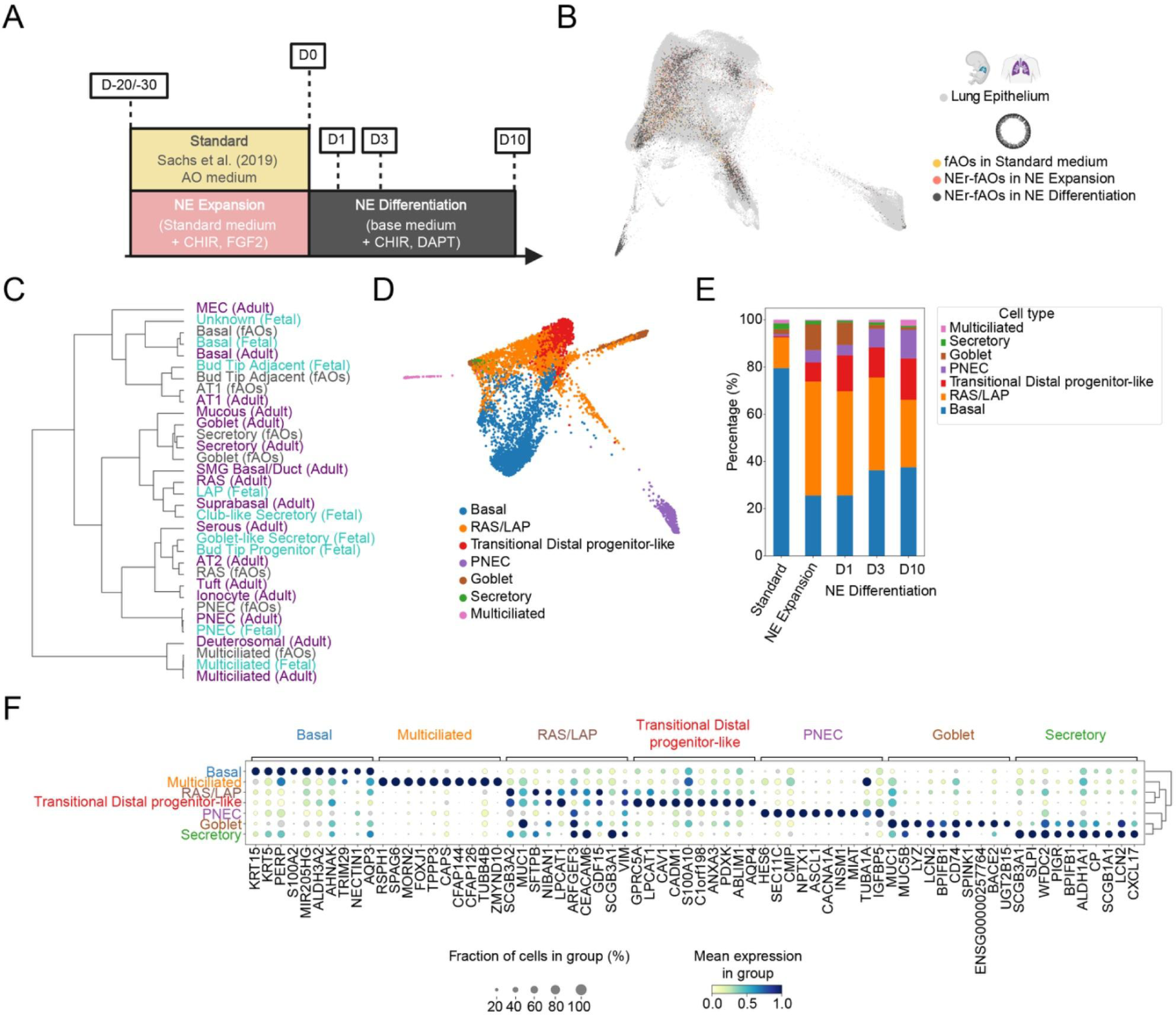
fAOs preserve a high degree of transcriptomic similarity with *in vivo* lung epithelial cells. (A) Schematic of scRNA-seq sampling from fAOs cultured in standard, and NEr-fAOs in NE Expansion, and NE Differentiation media across the differentiation time course. (B) FDL visualisation of fAOs scRNA-seq data integrated with *in vivo* lung epithelium scRNA-seq datasets (fetal and adult). (C) Hierarchical clustering of transcriptome between fAOs cells annotated by Celltypist (black) and *in vivo* cell type from Fetal (blue) and Adult (purple) Lung Epithelium. (D) FDL visualisation of fAOs scRNA-seq data colored by cell type (E) Stacked bar plot showing cell type proportion in fAOs growing in the different media conditions. (G) Dot plot of top 10 genes from each cell type in fAOs.

The PNEC cluster (363 total cells) contained 6 cells from fAOs in standard media, 50 cells from NEr-fAOs in NE expansion, and 313 cells from NEr-fAOs in differentiation media. As we had not observed the presence of PNECs in fAOs in standard media beyond passage 6, we reasoned this could be due to the early passage (P4 and P6) of the sequenced lines. To explore this further and assess PNEC identity, we calculated the PNEC signature score of cells from the PNEC cluster by the media dataset they came from, standard, NE Expansion, or NE Differentiation. This analysis showed that the PNEC Signature score of standard media-derived cells in the PNEC cluster was close to 0 and lower than for cells derived from NE Expansion and NE Differentiation, with NE Differentiation cells showing the highest PNEC Signature score (Figure S3i).

Further confirming the identities of cell types in fAOs and NEr-fAOs, analysis of differentially expressed genes across distinct cell types showed that well-known cell type markers were expressed in their respective categories (Fig. 3f), such as KRT5 for basal cells, LYZ for goblet cells, SCGB3A1 for club cells, FOXJ1 for multiciliated cells, SCGB3A2 for RAS/LAP cells, and ASCL1 for neuroendocrine cells. To enable interactive and user-friendly exploration of gene expression patterns in the dataset, we developed an online Shiny application, the NEr-fAOs scRNA-Seq Explorer (https://apps.embl.de/nerfaos_explorer/). Overall, our findings indicate that fAOs in standard media and NEr-fAOs in NE Expansion and NE Differentiation media contain a fetal progenitor population known as LAP cells, along with other airway epithelial cells that maintain a high degree of transcriptomic similarity to their *in vivo* counterparts.

### PNECs in NEr-fAOs resemble human primary PNECs

Having established that fAOs and NEr-fAOs maintain a high degree of transcriptomic similarity to *in vivo* lung epithelial cells, we next sought to compare PNECs in NEr-fAOs to PNECs derived from other published human airway models^14,19,21^ (Figure 4a), and to evaluate how closely *in vitro* PNECs resemble human primary adult or fetal PNECs^24,32^. PNECs derived from NEr-fAOs exhibited expression levels of canonical neuroendocrine markers - including *ASCL1*, *UCHL1*, *CHGA*, *GRP*, *CALCA*, *SCGN*, and *PCSK1N* - comparable to those observed in tissue-derived PNECs (Fig. 4b). While PNECs from other models also showed comparable expression of these canonical neuroendocrine markers, induced PNECs (iPNECs), 2D cultures derived from induced pluripotent stem cells following a 91-day air-liquid interface (ALI) differentiation protocol, showed markedly reduced expression of key markers compared to both tissue PNECs and PNECs from other in vitro systems (Fig. 4b).

**Figure 4.**
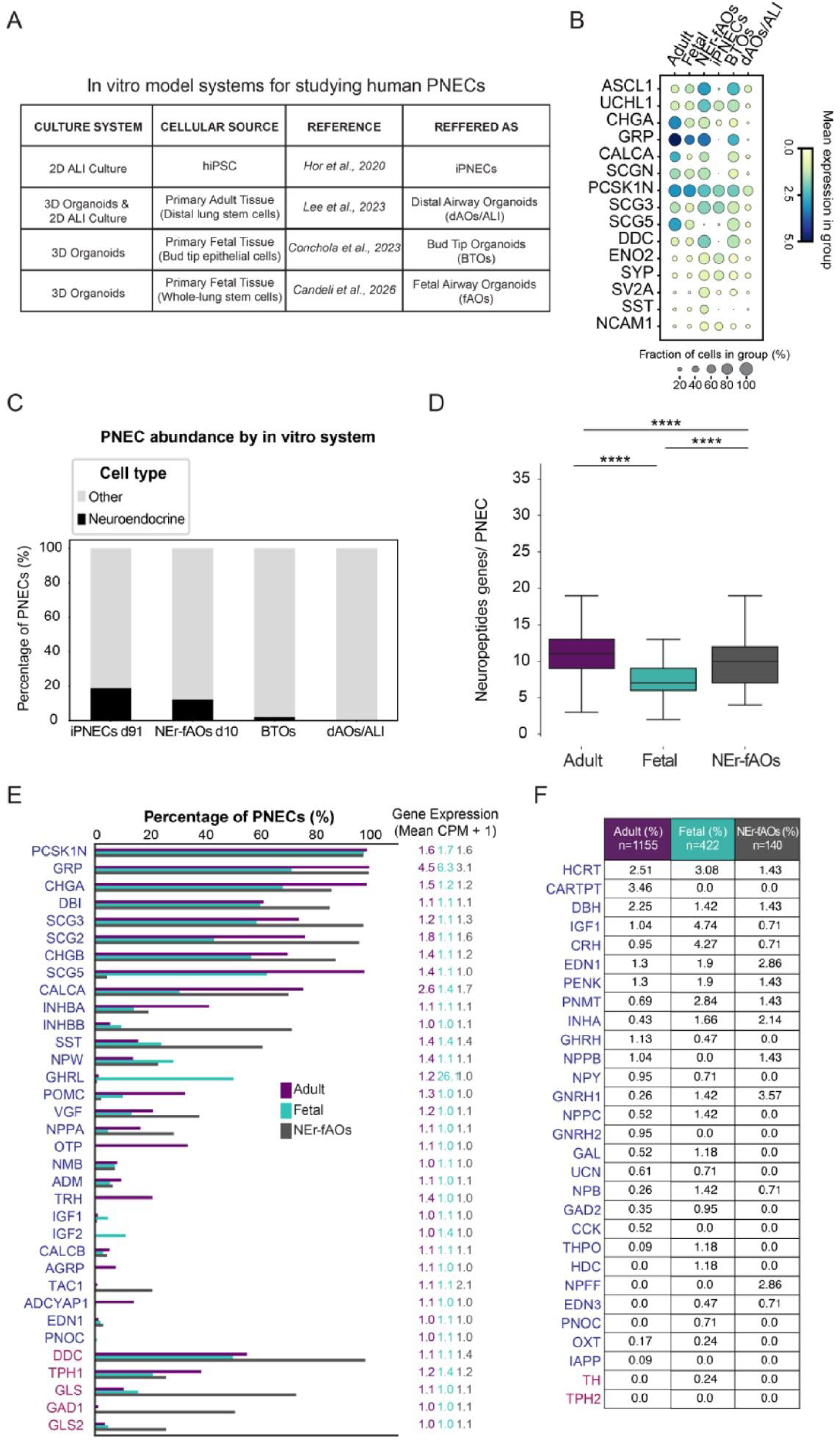
PNECs in Fetal Airway Organoids resemble human Fetal primary PNECs. (A) Table showing in vitro model systems for studying human PNECs. (B) Dot plot showing expression of canonical neuroendocrine markers across different in vitro model systems and in vivo lung epithelium. (C) Stacked bar plot showing yield of PNECs across different in vitro model systems. (D) Box plot showing the number of peptidergic genes expressed per PNEC in adult lung epithelium, fetal lung epithelium, and fAOs. (E) Bar plot representing the frequency of expression of individual peptidergic (blue) and neurotransmitter biosynthesis (red) genes in PNECs from adult lung, fetal lung, and fAOs. (F) Table comparing the expression of peptidergic (blue) and neurotransmitter biosynthesis (red) genes identified in rare PNECs (<3% prevalence) in primary lung epithelium with their expression in fAO-derived PNECs.

We next compared PNEC differentiation efficiency across in vitro systems (Fig. 4c). To mitigate potential biases introduced by inconsistent cell type annotations across different datasets, we performed automated PNEC identification in all datasets using CellTypist^31^, integrating previously published single-cell atlases of the human lung epithelium as reference datasets ^24,32^ (Fig. S4a). Among the systems examined, our 10 day differentiation protocol of NEr-fAOs in the NE Differentiation media yielded the second highest proportion of PNECs relative to total cells. From our day 10 dataset of NEr-fAO differentiation, we identified 140 PNECs, corresponding to 12% of all sequenced cells from that time point (140/1,158). By comparison, in the tissue datasets, PNECs constituted only 4% of fetal epithelial cells (422/10,614) and 0.85% of adult epithelial cells (1,155/135,863). The PNEC yield of NEr-fAOs was surpassed only by iPNECs generated via a 91-day ALI differentiation protocol (18%). In comparison, other models demonstrated substantially lower PNEC yields, including bud tip organoids (1.6%) and distal airway organoids (0.07%) (Fig. 4c).

PNECs act as multimodal sensors to perceive multiple airway stimuli and relay signals that either act locally or systemically to regulate homeostasis. The signals relayed by PNECs include neurotransmitters and neuropeptide and peptide hormones. To determine whether PNECs in NEr-fAOs retain the expression of neurotransmitter biogenesis genes, and neuropeptide and peptide hormone genes (“peptidergic” genes), we assessed their expression. NEr-fAO-derived PNECs expressed 48 peptidergic genes, with individual cells expressing 9.7 ± 3.9 (mean ± SD, range 4–19), comparable to adult (50 genes, 11.3 ± 3.2, range 3–24) and fetal PNECs (48 genes, 7.7 ± 3.5, range 0–34) (Fig. 4d and S4b, Table S2). Closely reflecting the diversity observed in primary fetal (391/422, 92%) and adult (948/1155, 82%) PNECs, a large proportion of NEr-fAO-derived PNECs (117/140, 83%) expressed unique combinations of peptidergic genes (Table S2).

Many peptidergic genes detected in only a small fraction of PNECs in the lung epithelium (<5%) also remain rare in NEr-fAO-derived PNECs, with the notable exception of TAC1—reported in lung neuroendocrine tumors^33^— which was notably enriched (31%) in NEr-fAOs (Fig. 4e and S4c). Expression patterns of other more commonly detected peptidergic genes were broadly conserved in fAOs, though some differences were observed—*GHRL*, for instance, was exclusive to fetal PNECs, while *OTP*, a key neurodevelopmental transcription factor ^34,35^, was exclusive to adult PNECs (Fig. 4e and S4c).

NEr-fAO-derived PNECs retained expression of biosynthetic genes for neurotransmitters serotonin (TPH1) and GABA (GAD1), the major neurotransmitters expressed by human PNECs (Fig. 4e and S4c). Expression of key glutamatergic (GLS, GLS2) genes was detected in both tissue-derived PNECs and NEr-fAO-derived PNECs. In contrast, expression of the dopaminergic gene TH, histaminergic gene HDC, catecholaminergic (DBH, PNMT) genes were detected in none or very few (<3%) PNECs across adult lung, fetal lung, and NEr-fAOs (Fig. 4f and S4d).

Despite the highest yield of neuroendocrine cells in the iPNEC systems, iPNECs show lowest diversity in the combination of peptidergic genes (iPNEC=517/786, 65%) and lower number of neuropeptides expressed per PNEC 5.7±4.3 (mean ± SD, range 0–34) (Fig. S4b and Table S2). Canonical markers such as *GRP, CHGA, CALCA,* and *SST* were markedly downregulated in iPNECs. Conversely, genes like *IGF2, SCG3, GAL, NPCC, PNMT,* and *TH* were significantly upregulated compared to both adult and fetal PNECs (Fig. S4c,d).

Overall, these results demonstrate that the NEr-fAO platform enables efficient generation PNECs at yields second only to the iPNEC system, while more closely recapitulating the transcriptomic features and molecular diversity in peptidergic gene expression of primary fetal and adult PNECs.

### Uncovering PNEC heterogeneity and differentiation mechanisms

The rarity of PNECs has made it challenging to investigate their molecular diversity and the mechanisms underlying their heterogeneity. To better characterize PNEC states and their relationship to differentiation processes, we performed clustering analysis of PNECs from our time-resolved scRNA-seq in NEr-fAOs under NE differentiation. For this subclustering analysis we extracted annotated PNECs (pertaining to leiden cluster 6) and of cells from leiden cluster 11 which showed high expression of the PNEC lineage transcription factor, *ASCL1*, suggesting it might represent a progenitor population undergoing neuroendocrine differentiation (Fig. S5a,b).

This sub-clustering analysis uncovered transcriptional heterogeneity within the PNEC population, resolving 5 discrete molecular states not previously defined, each characterized by distinct marker gene expression, including *NIBAN1*, *CALML3*, SST, *CALCA*, and *NEUROD1* (Fig. 5a,b), along with additional subcluster-specific upregulated genes (Fig. 5c, Table S3). Consistent with prior developmental atlases, which identified *MYCL*⁺ PNEC-progenitors and *GRP^high^* and *GHRL^high^* PNEC subsets in the developing human lung^36,37^, we observed correlated expression of *CALML3*, *CALCA*, and *NEUROD1* with *MYCL*, *GRP*, and *GHRL*, respectively, in fetal tissue-derived PNECs (Fig. 5d). Comparable correlated expression patterns were detected in NEr-fAO-derived PNECs, arguing that the *in vitro* PNEC states that we observe in NEr-fAOs are analogous to those seen in the endogenous tissue (Fig. 5e). The notable exception was *GHRL*, which was minimally expressed in NEr-fAOs (Fig. 5e,f).

**Figure 5.**
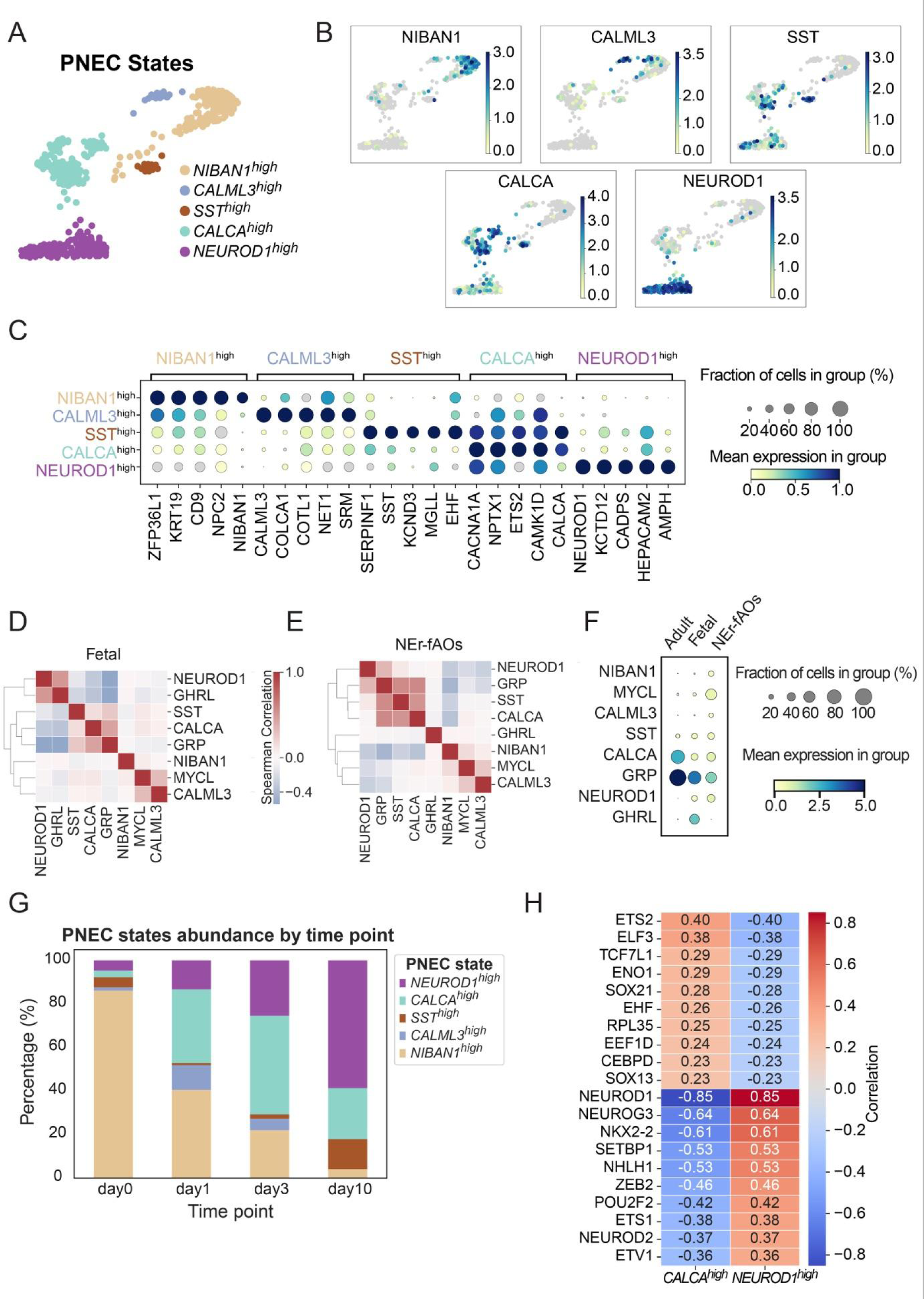
Early precursor and terminally differentiated NE cell states are captured in NEr-fAO organoids growing in NE differentiation media. (A) FDL visualisation of subsetted PNECs from fAOs colored by subcluster (PNEC state). (B) FDL visualisation of gene expression of PNEC state markers *NIBAN1*, *CALML3*, SST, *CALCA*, *NEUROD1* (C) Dot plot of top 5 genes from PNEC states. (D) Spearman correlation matrix showing pairwise gene expression correlations among PNEC state markers in PNECs from fetal lung epithelium and (E) NEr-fAOs. (F) Dot plot comparing expression of PNEC state marker genes in PNECs derived from *in vivo* adult and fetal airway epithelium, and derived from NEr-fAOs. (G) Stacked bar plot illustrating the proportions of different PNEC states in NEr-fAOs across various time points during NE differentiation. (H) Heatmap showing Cellrank-computed top 10 transcription factors driver for the NEUROD1*^high^* or CALCA*^high^* trajectories. The plot displays the correlation between fate probabilities toward the NEUROD1*^high^* or CALCA*^high^* states (Supplementary Figure F) and gene expression.

Interestingly, the *SST^high^*, *NEUROD1^high^*, and *CALCA^high^* PNEC states exhibited enrichment at day 10 of differentiation, suggesting they represent more differentiated PNECs. On the other hand, the *NIBAN1^high^* and *CALML3^high^* states show enrichment at early time points and depletion throughout differentiation, suggesting this population corresponds to precursor cells at early stages of PNEC differentiation (Fig. 5g and S5c).

To interrogate this finding using a different, but complementary, computational approach, we used CellRank. CellRank integrates directional information from per-cell velocity estimates with standard pseudotime inference based on cell-cell similarity to infer global transcriptional dynamics, allowing for the robust identification of initial, and terminal states as well as putative lineage drivers^38^. Applying CellRank to NEr-fAO PNECs, the *NEUROD1^high^* and *CALCA^high^* populations were consistently identified as terminal states, while the *NIBAN1^high^* populations were consistently defined as initial states (Fig. S5d-f).

To further elucidate the gene regulatory dynamics underlying PNEC differentiation, we leveraged CellRank to identify potential transcription factor (TF) drivers guiding differentiation toward either the *NEUROD1^high^* or *CALCA^high^* states (Fig. 5h and S5g). Through these analyses, we identified several known regulators of neuroendocrine differentiation in the intestine and pancreas, including *NEUROG3*, *NKX2-2,* and *NEUROD1* as drivers of the *NEUROD1^high^* state^39^. Drivers of the *CALCA^high^* state included genes associated with neuroendocrine and neuronal differentiation, and with some neuroendocrine cancers, such as *SOX21*, *ELF3*, and *EHF*^40–43^. Altogether, our work sheds light on the heterogeneity and differentiation processes of PNECs. Through these analyses we captured early and terminal states through dynamic PNEC differentiation in NEr-fAOs, and we uncovered putative gene regulators of these differentiation processes.

### NE Expansion media promotes a more distal airway identity

We were intrigued by the finding that simply adding CHIR and FGF2 to fAO media (NE Expansion media) enabled the generation of PNECs in NEr-fAOs from donor tissue that failed to do so when isolated and grown in standard media, which lacks these factors. To better understand this, we compared global gene expression changes in NEr-fAOs grown in NE Expansion versus fAOs grown in standard medium. This comparison revealed that cells from the two conditions occupied different territories within the force-directed graph embedding, indicating divergent transcriptomic landscapes (Fig. 6a). Differential gene expression analysis revealed significant upregulation of RAS/LAP-associated genes in NEr-fAOs grown in NE Expansion medium (Fig. 6b, Table S4), including *SCGB3A2*, *SFTPB*, and *RNASE1*^14^. The increased fraction of RAS/LAP cells in NEr-fAOs in NE Expansion relative to fAOs grown in standard media could explain this result. Alternatively, given *SCGB3A2* and *SFTPB* expression in adult lung is reportedly confined to distal airway cells^44–46^, we reasoned that NE Expansion might induce a distal airway program more broadly. To distinguish these possibilities, we asked whether *SCGB3A2* and *SFTPB* are upregulated across cell types in NE expansion compared to standard conditions (Fig. 6c). Consistent with a shift toward distal identity under NE Expansion conditions, NEr-fAO-derived cells cultured in NE expansion medium showed higher expression of *SFTPB* and *SCGB3A2* than matched cell types in fAOs cultured in standard media.

**Figure 6.**
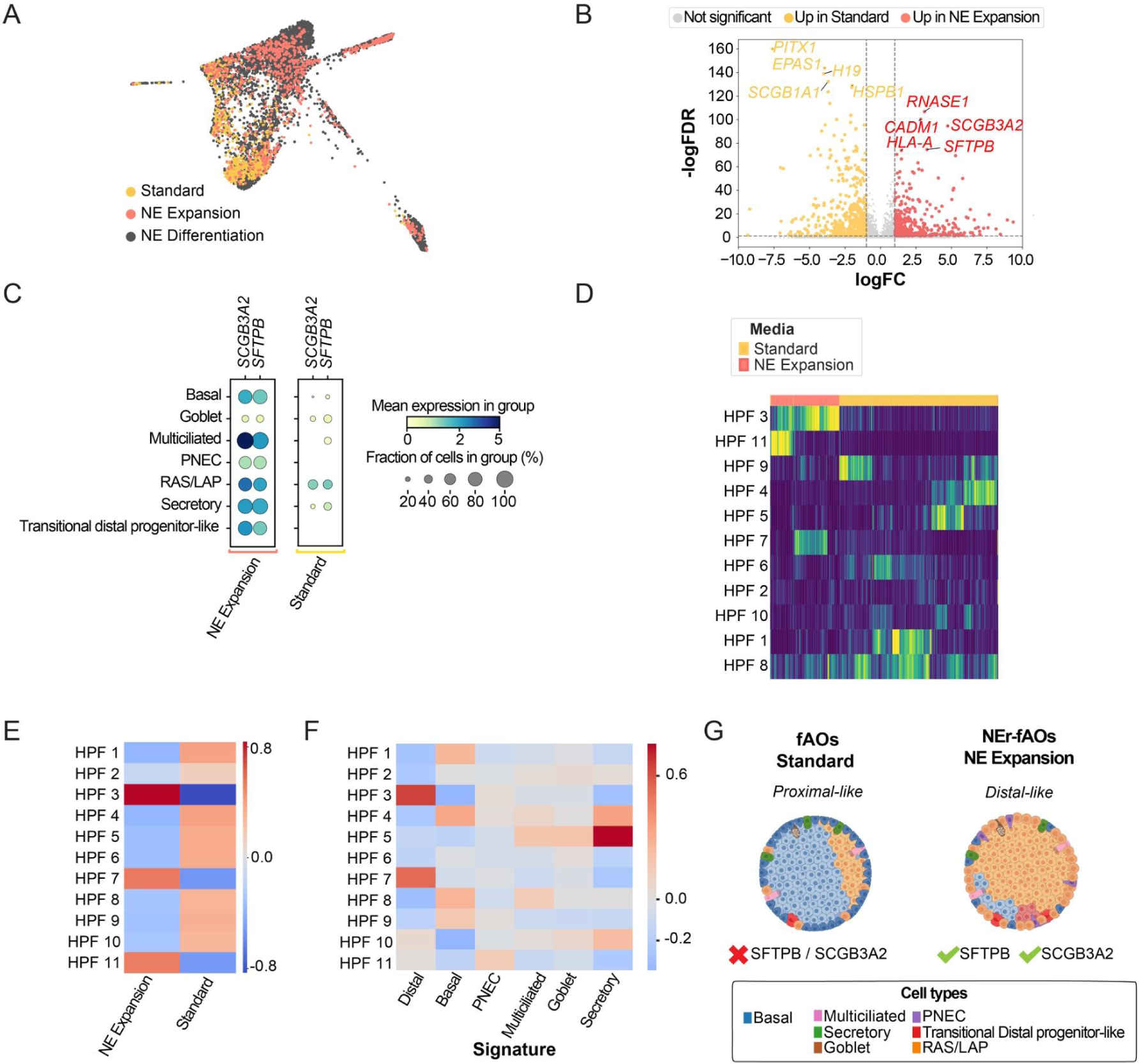
NE Expansion media promote distal airway identity. (A) FDL visualisation of scRNA-seq fAOs data colored by Media condition. Volcano plot of differentially expressed genes in fAOs grown in NE expansion (red) versus standard (yellow) medium (C) Dot plot of top distal genes *SCGB3A2*, *SFTPB* from each cell type in fAOs in standard or NE expansion media. (D) Hierarchical clustering of basal scHPF modules cell scores colored by media conditions. (E) Heatmap showing correlation of basal scHPF modules cell scores correlation with media condition (F) Heatmap showing correlation of basal scHPF modules cells scores with cell type signatures. (G) Schematic model of airway fate bias under different culture conditions. Cell types are indicated by color as shown.

To further interrogate this finding specifically in basal cells, a well-characterized airway cell type, we extracted and reclustered basal cells from NEr-fAOs in NE expansion media and fAOs in standard media. Basal cells from both media conditions expressed canonical basal cell markers such as, *KRT5*, *KRT15*, and *TP63*, but segregated within the force-directed graph embedding, indicating transcriptional divergence (Fig S6a, b). In line with our analysis of distal markers, basal cells from NEr-fAOs cultured in NE Expansion conditions displayed higher expression of these markers compared to fAOs grown in standard media (Fig. S6c).

To further interrogate comparative transcriptional dynamics within a specific cell type by an independent and complementary method, we applied the consensus single-cell Hierarchical Poisson Factorization (scHPF) algorithm^47^ across NEr-fAO-derived basal cells in NE expansion and basal cells derived from fAOs in standard media. Consensus scHPF identifies robust gene expression modules, representing additive transcriptional programs that contribute to variation across heterogeneous cell populations^47^. This approach identified 11 robust gene expression modules (scHPF factors) across these basal cells. Hierarchical clustering based on scHPF-derived cell scores for each gene expression module revealed stratification of basal cells according to media condition (Fig. 6d, Table S5). By correlating scHPF cell scores with culture conditions, we identified three modules (HPF3, HPF7, and HPF11) specifically enriched in NEr-fAO basal cells from NE Expansion medium (Fig. 6e). Consistent with our previous analysis and further suggesting that cells in NEr-fAOs have a more distal identity, cells corresponding to two of these modules (HPF3 and HPF7) are characterized by a high “distal score” (*SCGB3A2* and *SFTPB*) (Fig. 6f and S6d).

Collectively these data suggest a model where NE Expansion medium induces or selects for a more distal identity in NEr-fAO-derived cells and that this identity, which is enriched in RAS/LAP cells is permissive to subsequent PNEC differentiation (Fig. 6g). fAOs in standard medium appear to lack the capacity to support this distal identity and do not undergo downstream neuroendocrine lineage specification.

## DISCUSSION

Here we report NEr-fAOs, a versatile *in vitro* airway organoid platform derived from human fetal tissue stem cells (TSCs) that recapitulates key features of the airway epithelium and generates multiple differentiated cell types, including PNECs. Our results demonstrate that CHIR is sufficient to generate NEr-fAOs, which contain PNECs and RAS/LAP cells, the immediate precursors of PNECs^14^. Further, dual inhibition of GSK3 and NOTCH signaling cooperatively enhances and drives directed neuroendocrine differentiation, enabling robust enrichment of PNECs within NEr-fAO cultures.

Existing PSC-based approaches for modeling PNEC-associated diseases typically require prolonged differentiation protocols and have restricted capacity for long-term expansion. Prior TSC-derived airway models generally yield low PNEC abundance, limiting their applicability for mechanistic studies or disease modeling. In contrast, NEr-fAOs can be robustly expanded and efficiently differentiated into cultures enriched for PNECs while retaining diverse airway epithelial cell types. This combination of scalability and epithelial complexity provides a tractable experimental system for investigating the biology of this rare epithelial lineage in development and disease.

Through simultaneous inhibition of GSK3 and NOTCH signaling we can recapitulate PNEC differentiation in NEr-fAOs, capturing both progenitor and mature states that align with *in vivo* PNEC populations^36,37^. In future work, it will be interesting to explore how these pathways converge to promote the PNEC fate and whether similar logic can be leveraged to generate neuroendocrine lineages from other tissues such as intestine or pancreas.

Time-resolved scRNA-seq and lineage inference revealed that PNEC progenitors in NEr-fAOs give rise to distinct NEUROD1^high^ and CALCA-positive populations, where the latter includes *SST^high^* and *SST^low^* states. We identified transcriptionally similar populations in fetal lung tissue scRNA-seq datasets, supporting the idea that thse states represent functional specialization within the PNEC compartment. *CALCA*, which encodes calcitonin gene–related peptide (CGRP), is a neuropeptide involved in immune cell recruitment^7^ and airway repair following hypoxic injury^15^, suggesting that *CALCA^high^* PNECs may function as injury- or stress-responsive sensory transducers. In contrast, the *NEUROD1^high^*population expresses a transcription factor pivotal for neuronal differentiation^48^, suggesting a mature neuroendocrine state featuring enhanced neurosecretory and synaptic characteristics. This interpretation is consistent with developmental mouse studies implicating *NEUROD1* in PNEC subtype specification^49^. In the adult lung scRNA-seq datasets analyzed here, *NEUROD1* expression was exceedingly rare: among 1,155 PNECs, only one cell had detectable *NEUROD1*. Together, these findings support the idea that *NEUROD1^high^*PNECs are developmentally enriched, although platform-dependent sensitivity differences and scRNA-seq dropout remain plausible explanations for low detection of *NEUROD1* in adult datasets.

Notably, we did not detect GHRL-positive PNECs, a population characteristic of fetal development but also observed as a rare population in adult distal bronchioles^37,46^. Given reports that GHRL-positive PNECs rarely co-express *ASCL1* and that *NEUROD1* overexpression can drive differentiation into GHRL-positive PNECs in fetal lung organoids, it is likely that an extended differentiation period or temporal modulation of NOTCH inhibition (to reduce ASCL1 expression), may be required to access this in NEr-fAOs. Alignment of our NEr-fAO single cell transcriptomic dataset with fetal and adult datasets shows that NEr-fAO-derived PNECs closely resemble *in vivo* counterparts and express canonical neuroendocrine markers. Notably, ASCL1 and INSM1, key regulators of neuroendocrine differentiation^50,51^, were expressed at higher levels and in a greater proportion of NEr-fAO-derived PNECs than in PNECs from other *in vitro* systems, consistent with the enhanced differentiation achieved in our NE Differentiation media.

While PNECs generated in other systems also express canonical neuroendocrine markers at levels comparable to tissue, iPNECs, despite achieving the highest overall PNEC yield, showed reduced expression of these markers. While we cannot exclude the potential impact of cross-study detection differences, one potential explanation is the marked expression in this system of the mitogenic hormone, *IGF2*^52,53^, suggesting iPNECs might represent a proliferative PNEC state not typically observed in homeostasis.

NEr-fAO-derived PNECs retained expression of neurotransmitter biosynthetic genes (TPH1, GAD1, GLS, GLS2) and recapitulated the broad peptidergic diversity observed *in vivo*. While most peptidergic patterns were conserved, TAC1, reported in rare lung neuroendocrine tumors but absent in fetal and adult PNEC datasets, was uniquely detected in NEr-fAO-derived PNECs, suggesting our platform enables profiling of rare or transient neuroendocrine states. This highlights an important advantage of our NEr-fAO system. It will be interesting in future work, to explore how media conditions might impact PNEC peptidergic expression patterns.

Our observation that OTP, the neurodevelopmental transcription factor whose expression in tumors is associated with better prognosis in lung neuroendocrine tumors^54^, was detected in adult PNECs but not in fetal tissue or NEr-fAOs, is consistent with a late-stage differentiation program. Its loss in aggressive lung neuroendocrine tumors may reflect a reversion to a more immature, fetal-like transcriptional state. The absence of OTP-expression PNECs in NEr-fAOs may reflect either incomplete maturation in vitro or sampling limitations.

Our single-cell RNA-seq analyses highlight both the fidelity and utility of NEr-fAOs for research on PNEC biology. The NE differentiation protocol enabled high-depth SORT-seq of substantially more PNECs than is typically feasible from endogenous tissue, where PNECs are rare. Thus, NEr-fAOs enable robust transcriptomic profiling with far fewer sequenced cells than would be required from endogenous samples, providing an accessible source of PNECs for deep single-cell profiling and downstream mechanistic studies.

Beyond PNECs, our transcriptomic analysis suggests that NEr-fAOs are biased towards distal airway cell fates. This is supported by the prominence of RAS/LAP cells and basal stem cell heterogeneity with enrichment of the distal airway markers, *SCGB3A2* and *SFTPB*^45,55^. Consistent with their reported absence in fetal lung tissue, we did not detect adult distal basal cell populations, BC-1 and BC-2^55^, suggesting a broader distal signature rather than emergence of discrete basal identities. Because NE expansion medium differs from standard primarily by the addition of CHIR and FGF2, signalling downstream of these factors is likely driving this shift. Interestingly, CHIR is also required for directed PNEC differentiation.

Notably, the cell isolates that were used to establish fAOs in standard media and NEr-fAOs in NE expansion media were the same, highlighting the strong impact of media composition on the cell fates of the resulting organoids. In this case, NE expansion media composition could either be preferentially selecting for distal cell populations or it could be driving cells to a more distal state.

Recent studies highlight basal cell heterogeneity in both fetal^56^ and adult^55,57^ lung, reflecting variation along proximal-distal and apical-luminal axes. In the adult distal airways, basal cells in preterminal and terminal bronchioles (pre-TBs/TBs) have been proposed as progenitors of terminal airway secretory cells (TASCs)^57^. Related secretory progenitor populations, such as terminal respiratory bronchiole stem cells (TRB-SC) and respiratory airway secretory cells (RASCs), have also been implicated in distal airway and alveolar programs^55,58^. TASCs/RASCs/TRB-SCs share markers such as SCGB3A2 and SFTPB, which are also enriched in LAP cells, and our analyses suggest that RAS and LAP populations are analogous (Fig. 3). Consistent with this notion, the TASC/RASC/TRB-SC cell population is found in non-cartilaginous distal airways of the adult lung, the same anatomical niche where LAPs are found in the fetal lung. The shared molecular signatures and spatial localization suggest that TASC/RASC/TRB-SC may represent postnatal counterparts of fetal LAPs. It will be of interest to test whether adult distal airway progenitors can also generate PNECs and whether shared regulatory logic underlies these progenitor states.

In conclusion, NEr-fAOs provide a human fetal TSC-derived airway organoid platform for the long-term expansion and directed differentiation of a 3D pseudostratified airway epithelium enriched for PNECs. Systematic screening identified CHIR and FGF2 as key regulators of lineage commitment and revealed cooperative enhancement of PNEC differentiation by combined GSK3 and NOTCH inhibition. Single-cell transcriptomics showed that NEr-fAO-derived PNECs closely resemble their *in vivo* fetal and adult counterparts and capture developmental trajectories and molecular heterogeneity, including RAS/LAPs and distinct *NEUROD1*-positive and *CALCA*-positive subtypes. Together, these results establish NEr-fAOs as a tractable platform for dissecting human PNEC development and function, and studying neuroendocrine lineages in lung development, regeneration, and disease, including cancer.

## ACKNOWLEDGEMENTS

We thank Drs. Steve Lisgo and Nita Solanky and their teams at HDBR, including Jacqui Dobor and Berta Crespo Lopez for their invaluable support. We thank Single Cell Discoveries for SORT-seq single-cell sequencing services. We acknowledge financial support from Accelerate Lung Regeneration Consortium grant BREATH 12.0.18.002 of the Lung Foundation Netherlands (to H.C.), the European Molecualr Biology Lab (EMBL) core funding (to T.L.D.), an EMBO long-term fellowship (ALTF-21-2017) and a Marie Skłodowska-Curie IF grant 797966 – PNECtumor (to T.L.D.). N.C. was supported by a fellowship from the EMBL International PhD Program. A.M. was supported by a pilot grant from the EMBL Human Ecosystems Transversal Thene. A.G.G. was supported by a grant from the Associación Española Contrat el Cáncer (AECC, LABAE247362DAYT).

## AUTHOR CONTRIBUTIONS

T.L.D. and N.C. designed and conceived the study. T.L.D., L.dH., N.C., and A.M. generated the organoids. T.L.D., L.dH., F.S., N.C., A.M., and A.G.G. cultured and curated organoid lines and performed related experiments. N.C. processed and analyzed the sequencing data. T.L.D. and N.C. analyzed and interpreted the data and generated figures. J.L.M. contributed to study design. N.H. and J.S.V. contributed to analysis of sequencing data. J.S.V. developed the ShinyApp for scRNAseq data visualization. T.L.D. supervised the work. J.L.M. supervised N.H. T.L.D. directed the study.

## INCLUSION AND DIVERSITY

We support inclusive, diverse, and equitable conduct of research.

## FIGURES AND FIGURE LEGENDS

**Figure S1.**
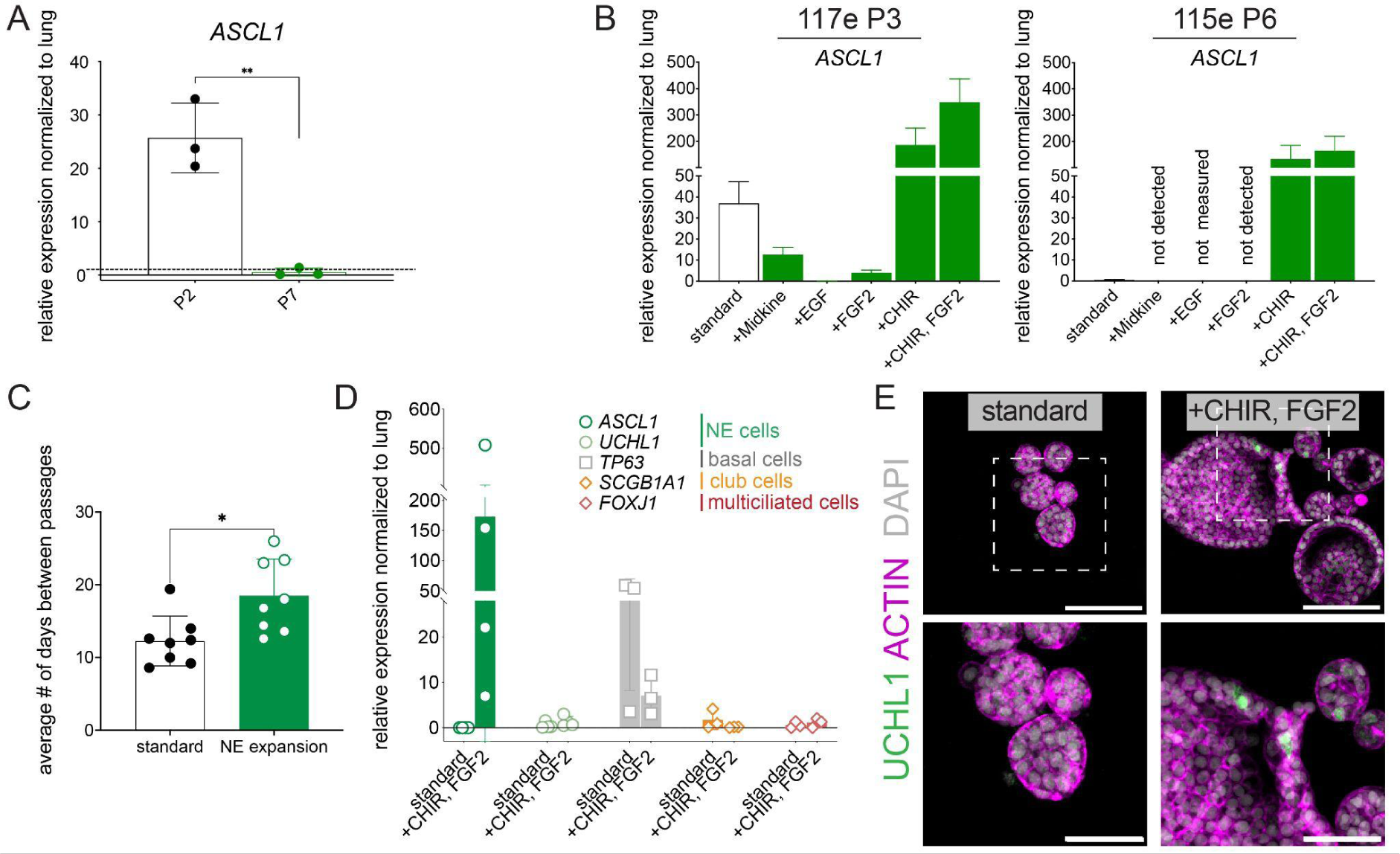
Optimised NE Expansion medium allows PNEC generation in fAOs. (A) qPCR analysis of PNEC (ASCL1) marker in fAOs cultured standard medium at early (P2) and late (P7) passage. Data correspond to 3 different lines. (B) qPCR analysis of PNEC (ASCL1) marker in fAOs cultured with indicated factors for 14 days. Data are represented as mean ± SD for 2 different lines. (C) Average passage time of fAOs grown in standard or NE Expansion medium. Data shown for 8 different lines. (D) qPCR analysis of PNEC (ASCL1) marker in fAOs at P9 cultured with indicated factors for 14 days. Data are represented as mean ± SD and correspond to 4 different lines. (E) IF image of fAOs in standard or CF2. ACTIN = membrane stain; UCHL1 = PNEC marker. Scale bars 100 μM for top panels and 50 μM for bottom panels.

**Figure S2.**
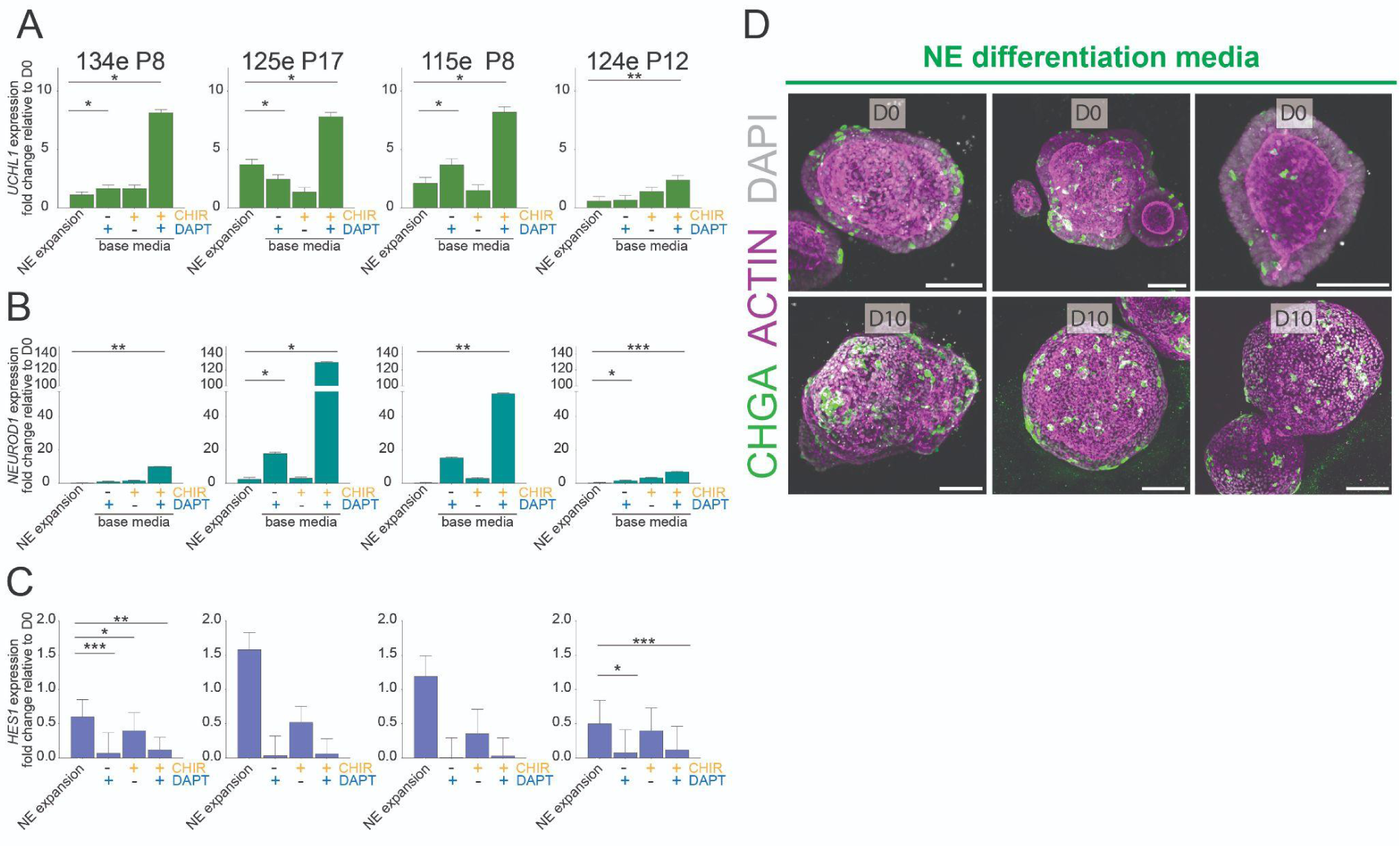
Dual inhibition of GSK3 and NOTCH signaling promotes PNEC differentiation. (A–B) qPCR analysis of early (*UCHL1*) and mature (*NEUROD1*) PNEC markers in NEr-fAO lines (n = 4) cultured in base medium supplemented with the indicated factors for 14 days. *P*-values using unpaired *t*-test are shown. (*) *P* < 0.05; (**) *P* < 0.01; (***) *P* < 0.001. (C) qPCR analysis of the NOTCH downstream effector *HES1* in NEr-fAO lines (n = 4) cultured under the same conditions. *P*-values using unpaired *t*-test are shown. (*) *P* < 0.05; (**) *P* < 0.01; (***) *P* < 0.001. (D) Immunofluorescence images of NEr-fAOs cultured in neuroendocrine differentiation media at day 0 and day 10. ACTIN = membrane stain; CHGA marks PNECs. Scale bars 100 μM.

**Figure S3.**
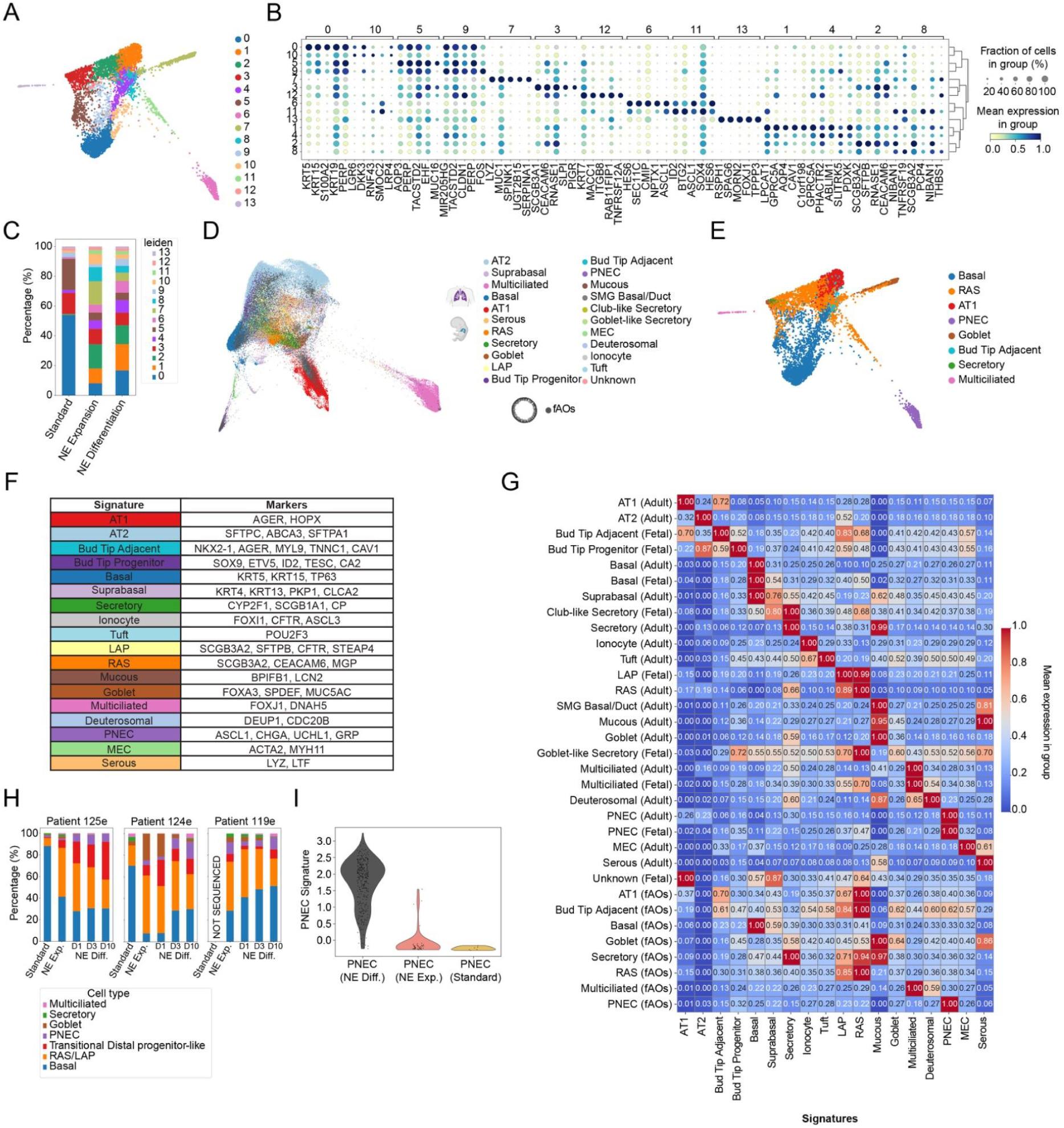
fAOs preserve a high degree of transcriptomic similarity with *in vivo* lung epithelial cells. (A) FDL visualization of overall Leiden clustering of fAOs cells. (B) Dot plot of top 5 genes from each Leiden cluster in fAOs. (C) Stacked bar plot showing Leiden clusters proportion in fAOs growing in the different media conditions. (D) FDL visualisation of fAOs scRNA-seq data integrated with *in vivo* lung epithelium scRNA-seq datasets (fetal and adult) colored by cell type. (E) FDL visualisation of fAOs cells colored by Celtypist-predicted cell types (F) Table of cell type signatures used for scoring using canonical markers for main lung epithelium cell types. (G) Matrix plot showing expression of cell type signature across fAOs and *in vivo* lung epithelial cell types. (H) Stacked bar plot showing cell type proportion in each of the 3 different sequenced fAO lines growing in the different media conditions. (I) Violin plot showing expression of the PNEC Signature across Celltypist annotated PNEC coming from different media conditions

**Figure S4.**
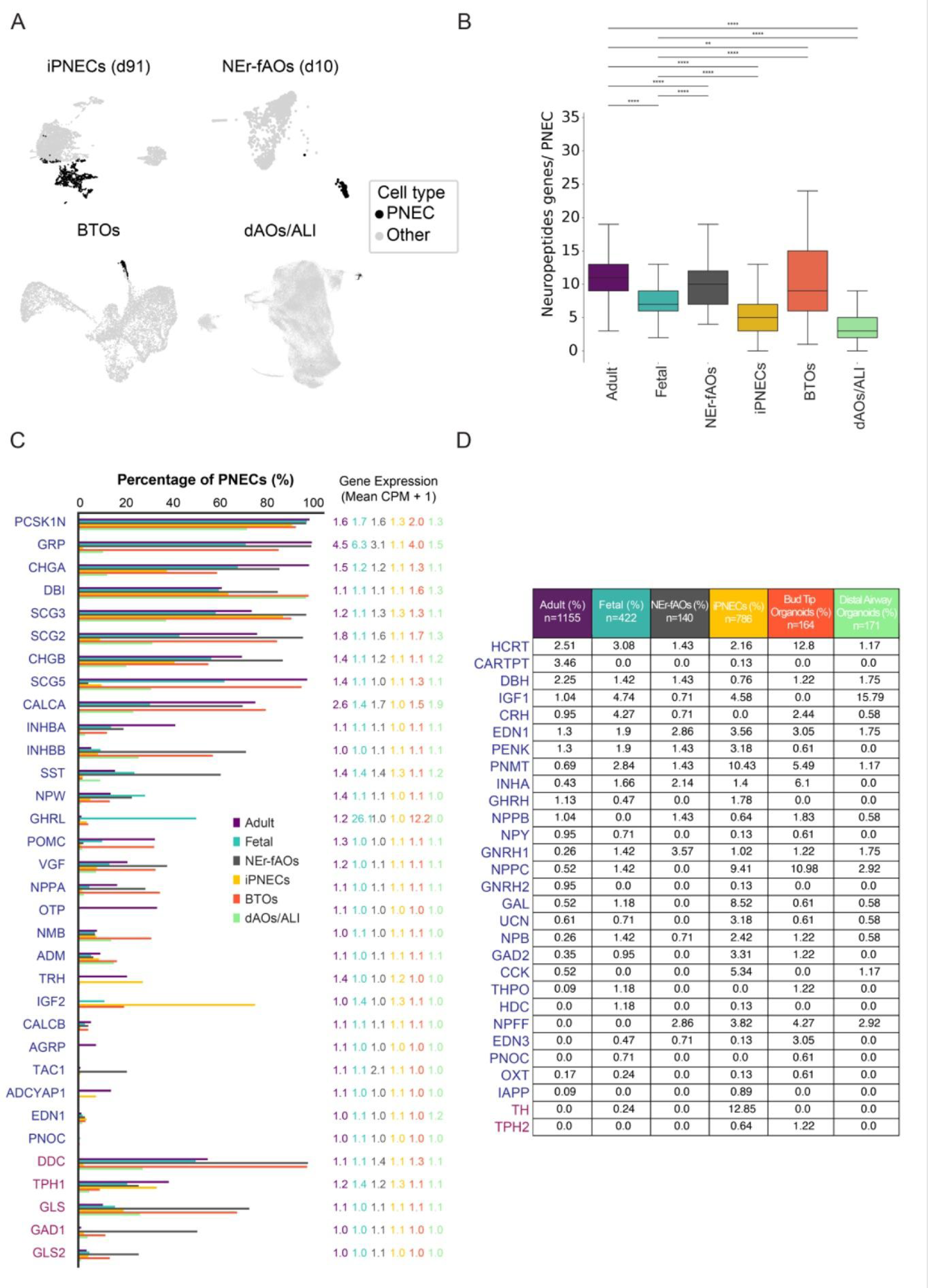
Characterisation of PNEC from *in vitro* model systems. (A) FDL visualisations of scRNA-seq datasets from in vitro model systems with PNEC (black) annotated by Celltypist. (B) Box plot showing the number of peptidergic genes expressed per PNEC in adult lung epithelium, fetal lung epithelium, and across different in vitro model systems. (D) Bar plot representing the frequency of expression of individual peptidergic (blue) and neurotransmitter biosynthesis (red) genes in PNECs from adult lung, fetal lung, and PNECs derived from different in vitro model systems. (C) Table comparing the expression of peptidergic (blue) and neurotransmitter biosynthesis (red) genes identified in rare PNECs (<3% prevalence) in primary lung epithelium with their expression in PNECs derived from different in vitro model systems.

**Figure S5.**
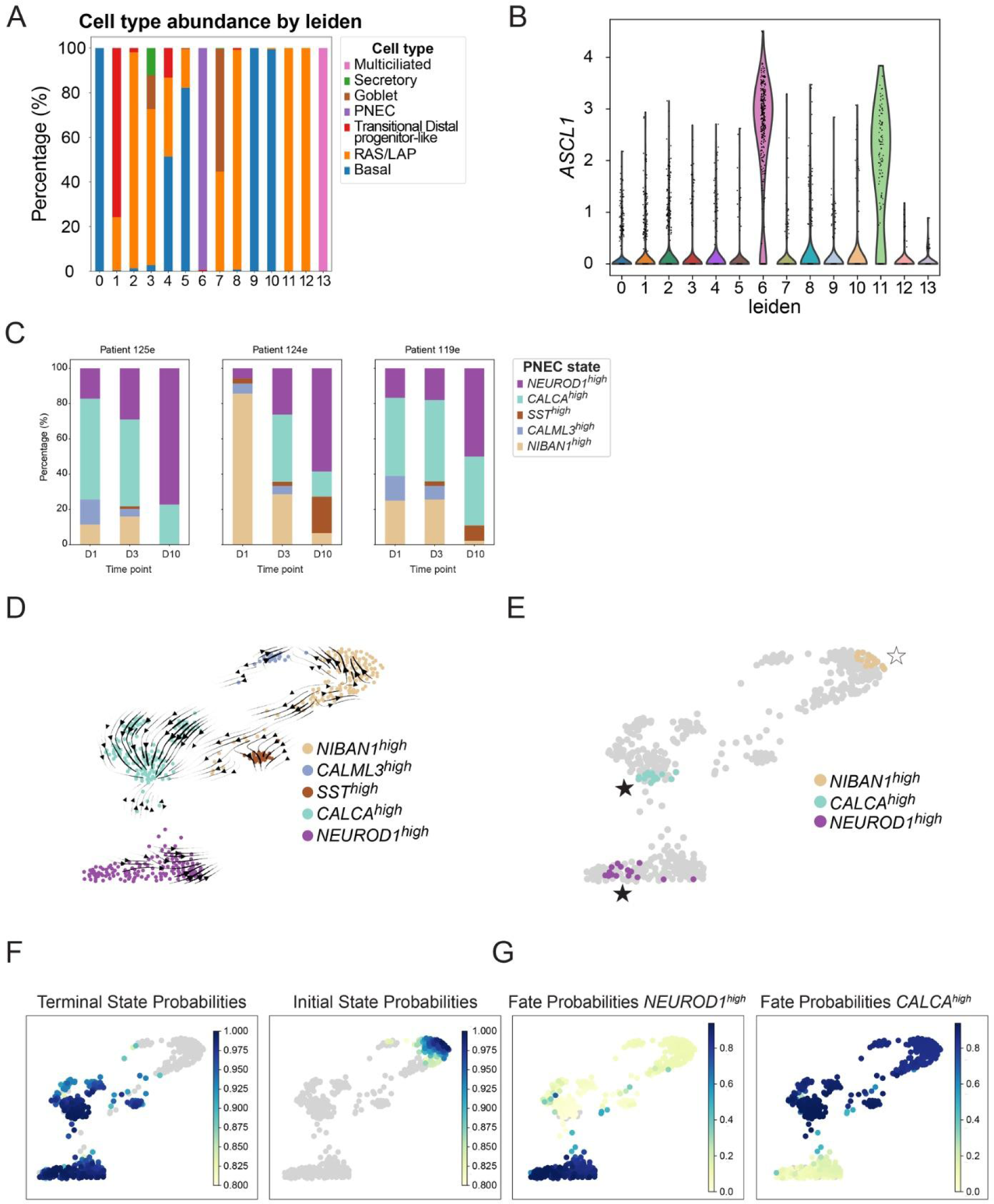
Identification of early precursor and terminally differentiated NE cell states. (A) Stacked bar plot illustrating the cell type proportions in fAOs across Leiden clusters. (B) Violin plot showing expression of the neuroendocrine canonical marker *ASCL1* across Leiden clusters. (C) Stacked bar plot illustrating the proportions of different PNEC states in NEr-fAOs from different donors and across various time points during NE differentiation. (D) FDL visualisation of PNEC states from NEr-fAOs colored by PNEC state showing RNA velocity vectors. (E) FDL visualisation of PNEC states showing initial (white stars) and terminal states (white star) imputed by CellRank. (F) CellRank prediction of initial and terminal state showing respective probabilities in the FDL visualisation. (G) CellRank prediction of differentiation fate probabilities towards the Neuroendocrine NEUROD1 ^high^ and CALCA ^high^ state.

**Figure S6.**
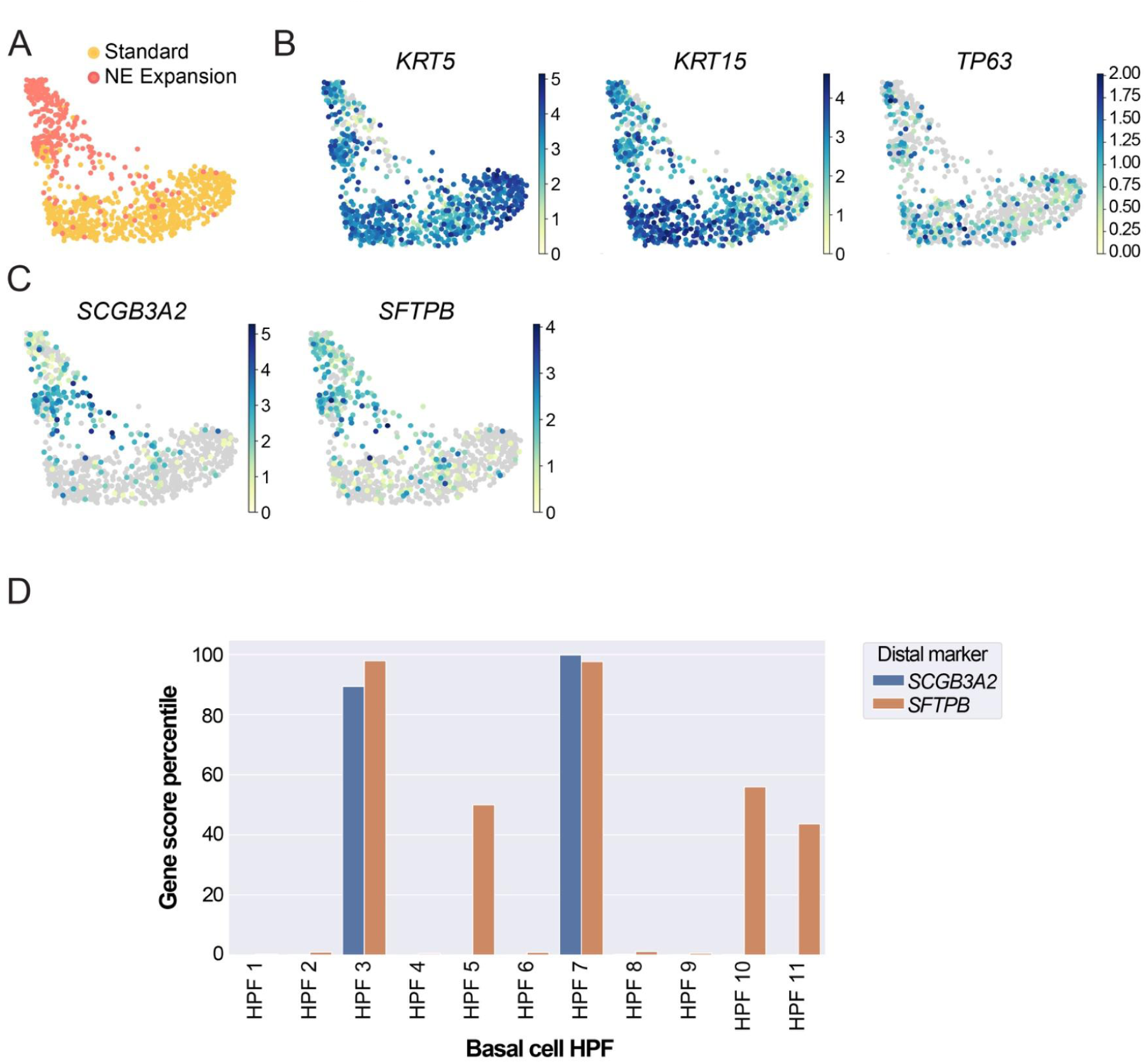
NE expansion promotes distal basal cell identity. (A) FDL visualisation of Basal cell extraction from scRNA-seq fAOs data colored by Media condition. (B) FDL visualisation of Basal cell extraction showing expression of Basal canonical markers KRT5, KRT15, TP63 (C) FDL visualisation of Basal cell extraction showing expression of LAP canonical markers SCGB3A2, SFTPB, CFTR, STEAP4. (D) Bar graph showing the distribution of distal airway genes (SCGB3A2, SFTPB) rank percentile across basal scHPF modules, indicating its relative activity level within each basal scHPF module.

**Table S1. List of lines generated and used in this study**

**Table S2.** Differentially expressed genes in fAO and NEr-fAO leiden clusters.

|  | Adult<br>(n=1155) | Fetal<br>(n=422) | NEr-fAOs<br>(n=140) | iPNECs<br>(n=786) | BTOs<br>(n=164) | dAOs/ALI<br>(n=171) |
| --- | --- | --- | --- | --- | --- | --- |
| Number of Neuropeptides Detected | 50/50 (100%) | 48/50 (96%) | 46/50 (74%) | 50/50 (92%) | 48/50 (96%) | 43/50 (86%) |
| Distinct Neuropeptide Expression Profiles per PNEC | 948/1155 (82%) | 391/422 (92%) | 117/140 (83%) | 517/786 (65%) | 150/164 (91%) | 100/171 (58%) |
| Average Number of Neuropeptides per PNEC (mean ± SD) | 11.3±3.2 | 7.7±3.5 | 9.7±3.3 | 5.7±4.3 | 10.3±5.6 | 3.6±2.8 |
| Range of Neuropeptides per PNEC | 3-24 | 0-34 | 4-19 | 0-34 | 1-24 | 0-16 |

**Table S3. Details about number of neuropeptide genes and neuropeptide profiles expressed in pulmonary neuroendocrine cells across *in vivo* lung tissue and in vitro model systems**

**Table S4. Differentially expressed genes in NEr-fAO PNEC subclusters**

**Table S5. Differentially expressed genes between NEr-fAOs in NE expansion and fAOs in standard media**

**Table S6. Genes corresponding to HPF modules in basal cells METHODS**

## KEY RESOURCES TABLE

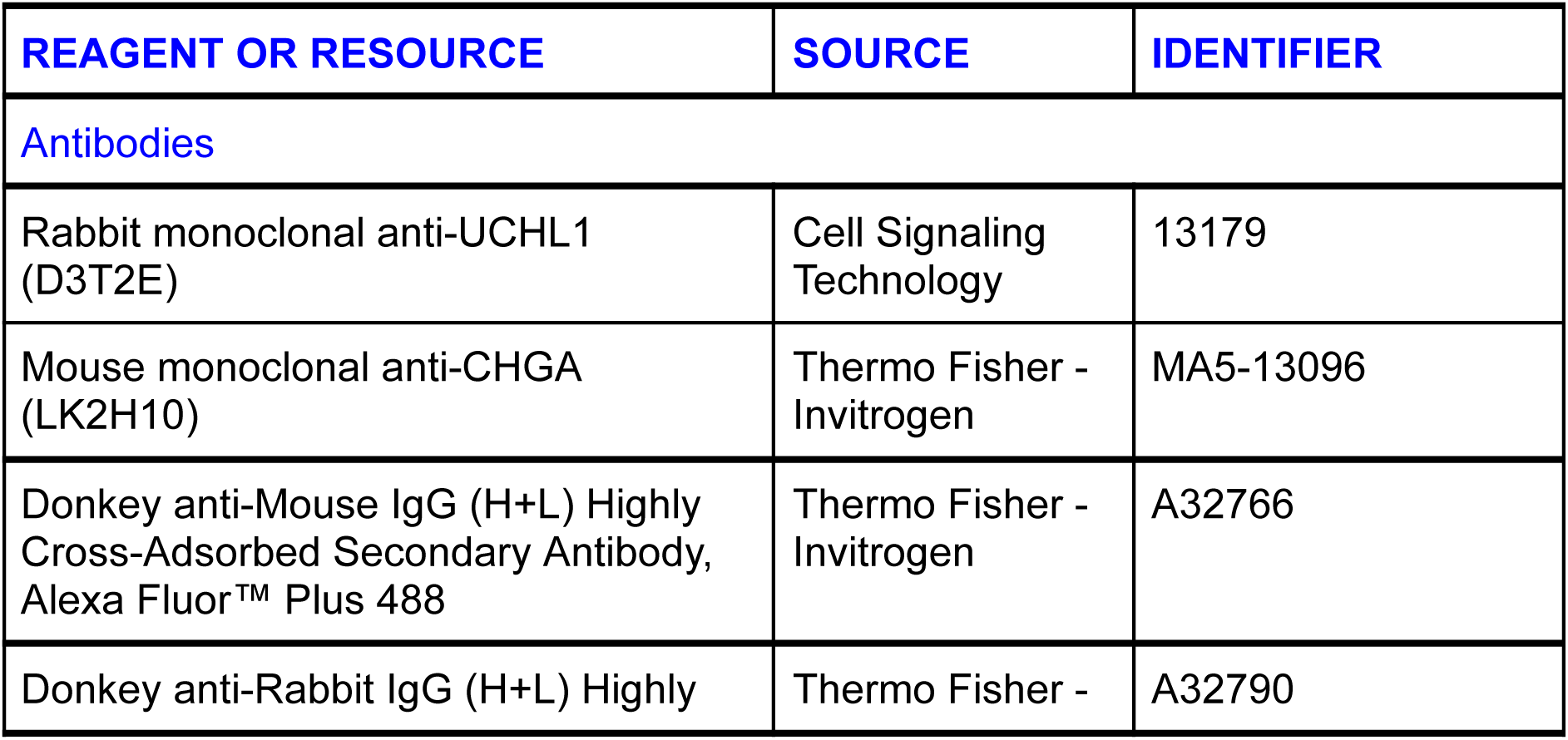
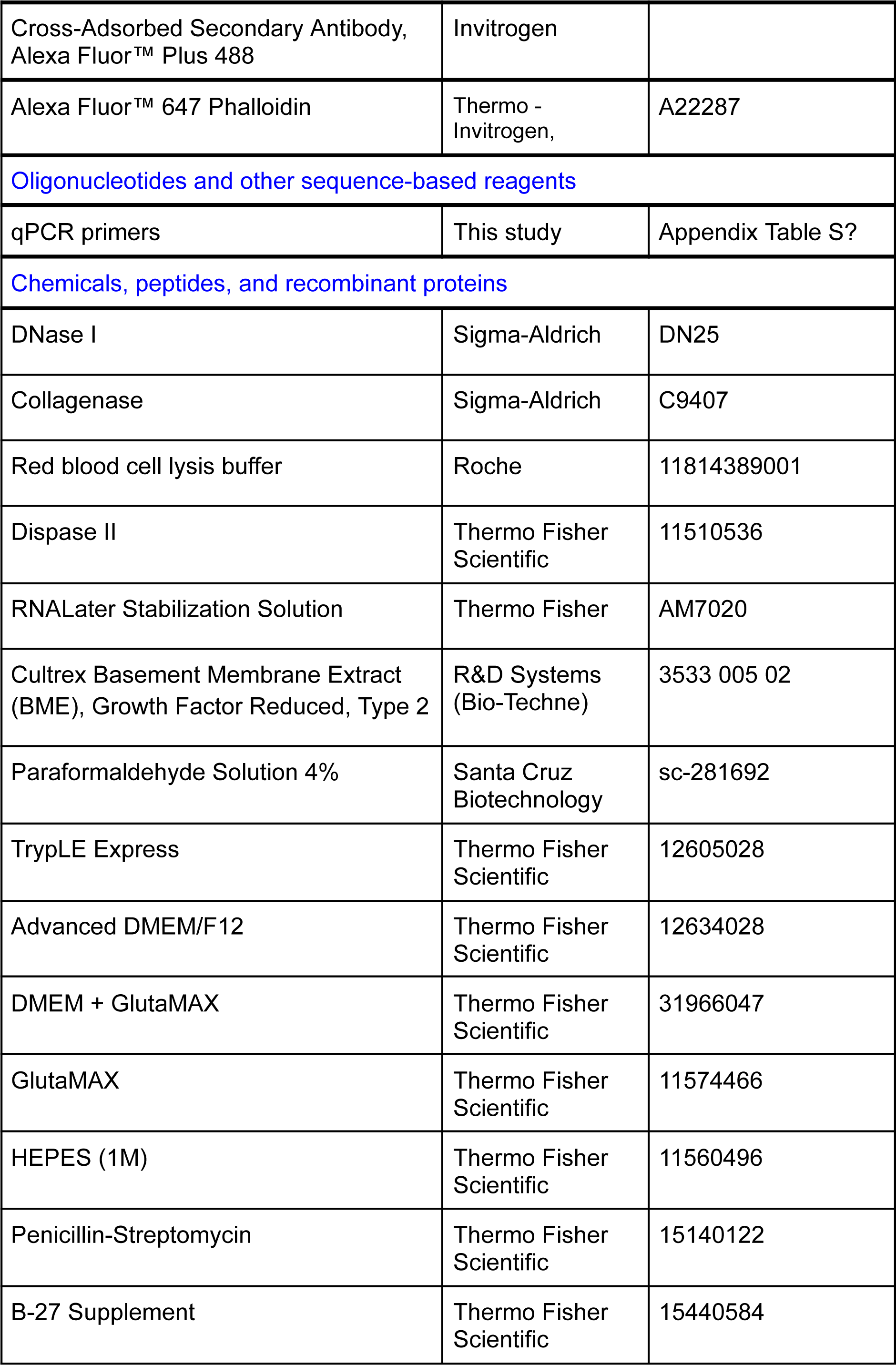
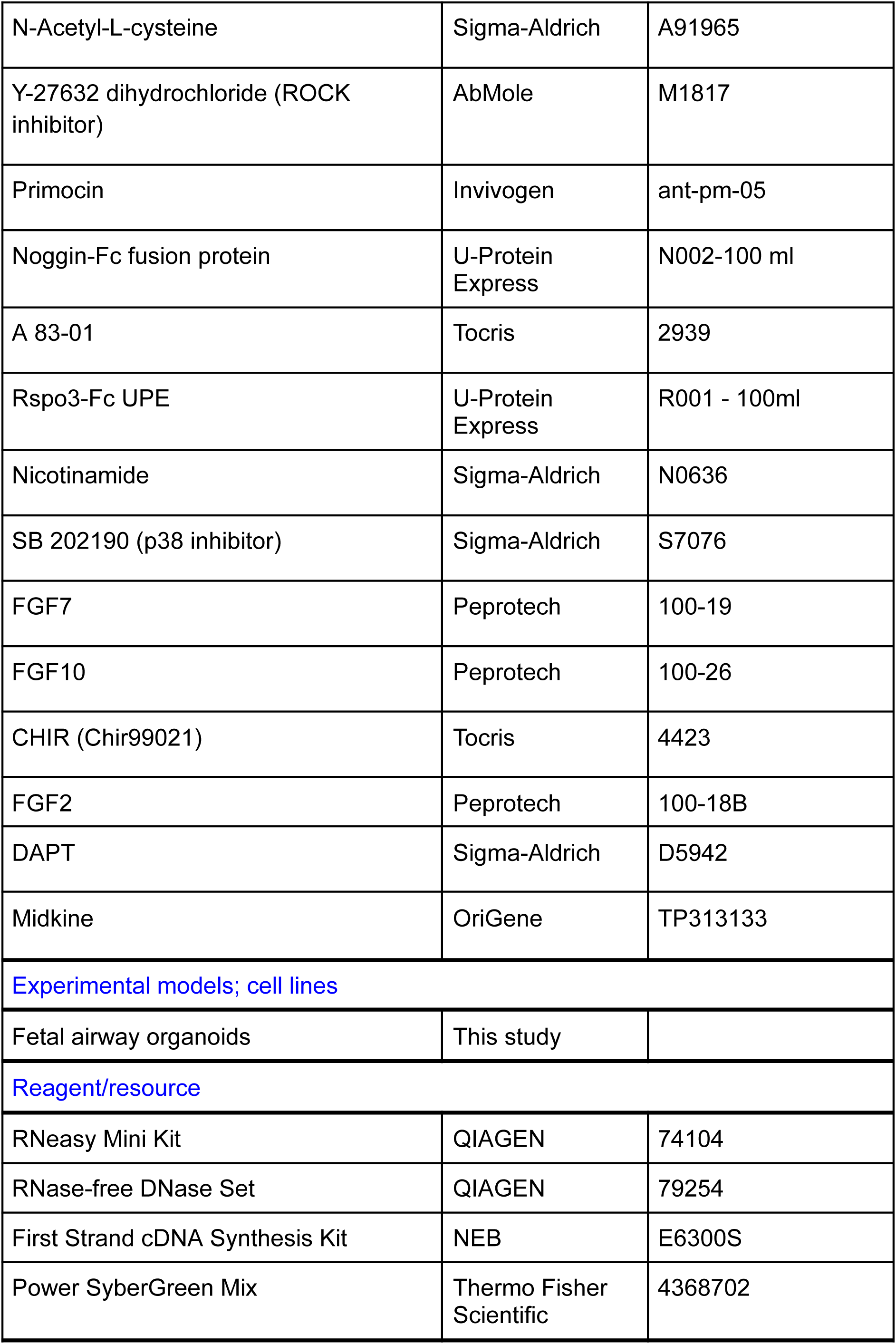
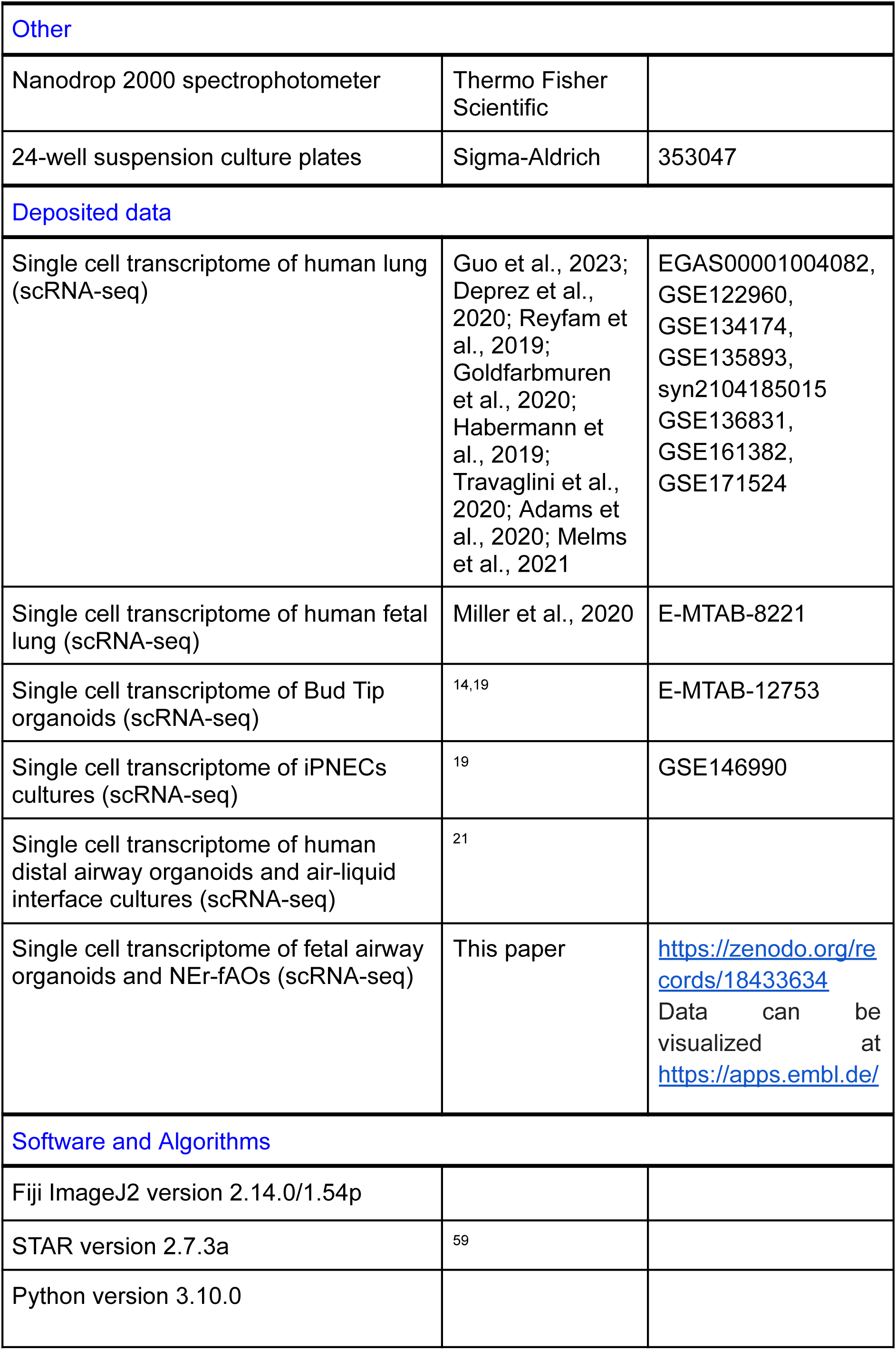
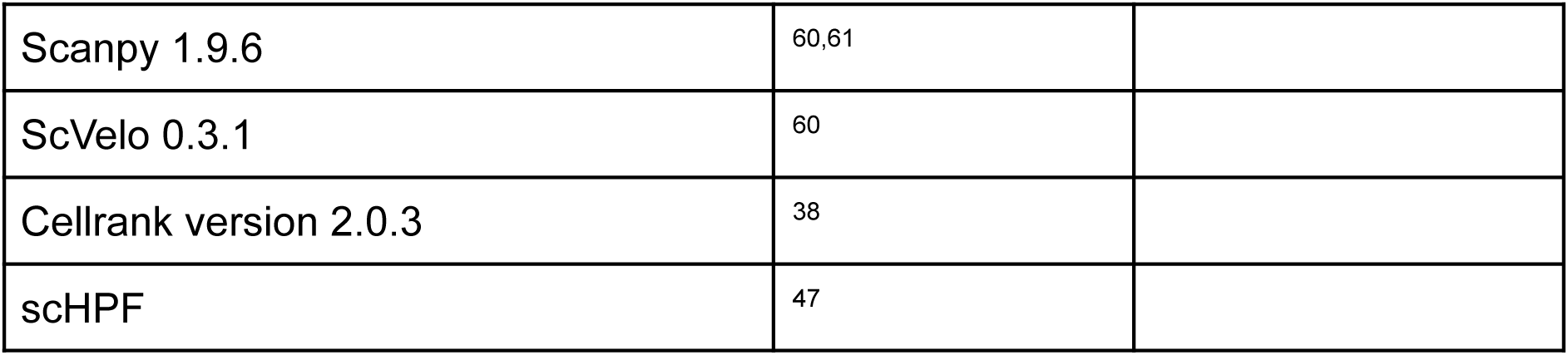

## RESOURCE AVAILABILITY

### Lead contact

Further information and requests for resources and reagents should be directed to and will be fulfilled by the Lead Contact Talya Dayton.

### Data and code availability

Count matrices and metadata corresponding to single-cell RNA sequencing data of this study have been deposited at Zenodo as https://zenodo.org/records/18433634 and are publicly available as of the date of publication. Due to donor privacy concerns, associated raw sequencing data (FASTQs) will be made available through controlled access from EGA. All bioinformatic processing pipelines are open-source and publicly available at https://github.com/daytonlab/NEr-fAOs_2026. The analysis scripts used to process the sequencing data and generate the associated figures are also available from the same GitHub repository.

In addition, an on-line ShinyApp tool has been established to allow interactive, user-friendly visualizations of gene expression in each population (https://apps.embl.de/nerfaos_explorer/).

## METHOD DETAILS

### Approval of studies involving human and patient-informed consent

Human fetal lung tissues were obtained from elective abortions from Leiden University Medical Center or the Human Developmental Biology Resource, under informed consent and ethical approval. Ethical approval for the use of human fetal tissue was provided by the Commission of Medical Ethics of Leiden University Medical Center (Leiden, NL), Human Developmental Biology Resource (HDBR, Cambridge and Newcastle, UK), and the Bioethics Internal Advisory Committee (BIAC) of the European Molecular Biology Lab (EMBL). In total, n = 12 tissues were used in this study. Tissues were from both sexes and ranged in developmental stage from 14 to 18 post conception weeks.

### Fetal Airway Organoids culture

#### Establishment of organoid lines

Solid lung tissue was transferred to a sterile 10 cm dish and minced using a disposable scalpel into small fragments (approximately 1–2 mm³). The minced tissue was resuspended in 10 mL of wash medium (DMEM containing 1x GlutaMAX (Thermo Fisher Scientific, 31966047), and 100 U/mL Penicillin-Streptomycin (Thermo Fisher Scientific, 15140122) and transferred to a 15 mL conical tube. The sample was pipetted up and down several times to mechanically dissociate the tissue further. 200 μL per sample of collagenase solution (20 mg/mL in AdDF++: Advanced DMEM/F12 (Thermo Fisher Scientific, 12634028) containing 1x Glutamax (Thermo Fisher Scientific, 11574466), 10 mM HEPES (Thermo Fisher Scientific, 11560496), and 100 U/mL Penicillin-Streptomycin) was added to the suspension, and the tube was tightly capped, sealed with parafilm, and incubated at 37 °C with shaking for 25 min. Following enzymatic digestion, 50 μL of DNase I (Sigma-Aldrich, DN25) were added, and the mixture was pipetted up and down to aid further dissociation. The digested suspension was filtered through a 100 μm cell strainer, using the plunger of a 3 mL syringe to facilitate passage of cells through the filter. The filter was subsequently washed with cold wash medium to recover residual cells. The flow-through was collected and centrifuged at 400×g for 5 min at 4 °C. The supernatant was aspirated carefully, leaving the cell pellet intact. The pellet was resuspended in 0.2–1mL Red Blood Cell Lysis Buffer (Roche, 11814389001) and incubated at room temperature for 5 min. The reaction was quenched by adding wash medium to a final volume of 6 mL, followed by centrifugation at 400×g for 5 min. The resulting white pellet was washed once with cold wash medium. The cell pellet was resuspended in ice-cold Cultrex growth factor reduced BME type 2 (Bio-Techne, 3533 005 02) and plated as 25–50 μL domes on 24-well suspension culture plates (Sigma-Aldrich, 353047). After BME polymerization (10–15 min at 37°C), the appropriate organoid culture medium was added.

#### Maintenance organoid culture

Cultures were maintained at 37 °C in a humidified incubator with 5% CO₂. Medium was replaced every 6–7 days, and organoids were passaged every 2-3 weeks: BME was dissolved in 1–2 mL of TrypLE Express (Thermo Fisher Scientific, 12605010) and transferred to a 15 mL conical tube. Samples were incubated in a 37 °C water bath for 2-5 min, followed by mechanical shearing through a serological pipette fitted with a P10 tip to increase shear force. Dissociation was monitored microscopically and stopped once a mixture of single cells and small clusters was observed. Following dissociation, 5–10 mL of wash medium was added to neutralize TrypLE. The suspension was centrifuged at 300 × g for 3 min at 4 °C. The supernatant was aspirated, and the pellet was resuspended in cold wash medium (approximately 5 μL per intended 24-well equivalent). Followed the addition of ice-cold BME (wash medium:BME ratio of ∼1:8), the suspension was gently pipetted to ensure uniform mixing, and cells were reseeded as above as at ratios (1:1– 1:6), depending on growth rate and density, allowing the formation of new organoids.

#### NE differentiation experiments

For the directed NE differentiation, NEr-fAOs were cultured in NE Expansion media. At 14 to 22 days of expansion, NE Expansion medium was removed and NE differentiation medium was added. Medium, including NE differentiation medium, was refreshed every 7 days.

#### Cryopreservation and Recovery

Organoids were cryopreserved as intact structures in FBS containing 10% DMSO, without enzymatic dissociation. For recovery, frozen vials were rapidly thawed, and organoids were transferred directly into BME domes in a 12-well plate. Cultures were allowed to recover for 2–3 days before further expansion or passaging.

### Culture Media, Growth Factors and Small Molecules

All experiments utilized a basal medium. Briefly, the basal medium consists of AdDF+++ (Advanced DMEM/F12 containing 1X Glutamax, 10 mM HEPES, and 100 U/mL Penicillin-Streptomycin) supplemented with 1X B27 supplement minus vitamin A (Fisher Scientific cat. no. 15440584), N-acetyl-L-cysteine (1.25 mM; Sigma-Aldrich, cat. no. A91965), Y-27632 dihydrochloride (ROCK inhibitor) (2.5 μM; AbMole, cat. no. M1817), and Primocin (50 μg/mL; InvivoGen, cat. no. ant-pm-05). The standard medium has been previously described^23^ and used the same basal medium, but was further supplemented with following growth factors and small molecules: Noggin-Fc fusion protein (1%; ImmunePrecise, cat. no. N002-100ML), A 83-01 (500nM ; Tocris, cat. No. 2939), R-spondin 3-Fc (Rspo3-Fc) (1%; ImmunePrecise, cat. no. R001-100ML), Nicotinamide (5 mM; Sigma-Aldrich, cat. no. N0636), SB202190 (p38 MAPK inhibitor) (1 μM; Sigma-Aldrich, cat. no. S7076), Fibroblast Growth Factor 7 (FGF7) (25 ng/mL; PeproTech, cat. no. 100-19), Fibroblast Growth Factor 10 (FGF10) (100 ng/mL; PeproTech, cat. no. 100-26). NE expansion medium consists of the standard medium, supplemented with CHIR99021 (3 μM; Tocris Bioscience, cat. no. 4423), and Fibroblast Growth Factor 2 (FGF2) (100 ng/mL; PeproTech, cat. no. 100-18B) added immediately prior to use. NE differentiation medium consists of the basal medium (supplemented with CHIR99021 (3 μM; Tocris, cat. no. 4423), and DAPT (γ-secretase inhibitor) (10 μM; Sigma-Aldrich, cat. no. D5942) added immediately prior to use. Other growth factors and small molecules were used at the following concentrations: Midkine (50 ng/mL; OriGene, cat. no. TP313133).

### Whole Mount Immunofluorescence on Organoids

Before immunofluorescence analysis cultured domes containing lung organoids were mechanically dissociated in ice-cold PBS. Released organoides were washed once with PBS and then fixed overnight in PBS with 4% paraformaldehyde (Santa Cruz, sc-281692) at 4°C. After fixation, organoids were permeabilized in PBS with 0.1% Tween-20 (Sigma, P7949) (PBST) for 30 min and separated into wells of a glass bottom 24-well plate (CellVis, P24-1.5H-N) for different staining. Afterwards, organoids were incubated with a blocking solution containing 5% Donkey Serum (Merck, 29663) in the wash buffer (PBS with 1% Triton™ X-100 (Fisher, BP151100) and 2g of BSA (Merck, 03116956001)) for 1h at 4°C. Next, organoids were incubated with primary antibodies diluted in the wash buffer overnight at 4°C. Then, organoids were washed 5 times with the wash buffer for 3 mins and subsequently incubated with diluted secondary antibodies and phalloidin for 3 hours at room temperature. Afterwards, another 5 washes were performed. Nuclei were then stained by incubating the organoids with 4,6-diamidino-2-phenylindole (DAPI; 1:2000, Thermo - Invitrogen, D3571) diluted in the wash buffer for 30 mins at room temperature (RT). Prior to imaging, organoids were washed 3 times and resuspended into the clearing solution consisting of 80% Glycerol (Thermo - Invitrogen, 17904) in PBST.

After at least 1 hour incubation in the clearing solution, images of stained organoids were acquired by using the Opera Phenix™ microscope. Settings were set to acquire in confocal mode and a binning of 2 with a 20X water-immersion objective. Z-stack images were collected as tiled volumes with a thickness of 350 µm and an xyz voxel size of 0.58 × 0.58 × 1 µm. All raw confocal microscope images were processed with the Fiji ImageJ2 software using published <u>macro scripts</u>. For optimal image visualization, Z-stacks were projected onto a single image where the background signal was removed, and brightness and contrast were enhanced.

### RNA Extraction and qRT-PCR Analysis

At least 1 well, containing 20-50 organoids, for each biological replicate was collected and RNA was extracted for QRT-PCR analysis. RNA was isolated from organoids using the RNeasy Mini Kit (QIAGEN, Cat# 74104) following the manufacturer’s instructions including DNaseI treatment (QIAGEN, Cat# 79254). RNA quality and concentration was determined on a Nanodrop 2000 spectrophotometer (Thermo Scientific). 1 ug of RNA for each sample was used to generate a cDNA library using the ProtoScript® First Strand cDNA Synthesis Kit (NEB, cat. No. E6300S). QRT-PCR was performed in triplicate on a Step One Plus Real-Time PCR System (Life technologies) using Power SyberGreen Mix (Thermo Fisher Scientific, cat. No. 4368702). Gene expression was quantified using the DDCt method and normalized by ACTB using the primers listed in the table below. Some data were plotted as a fold change of arbitrary expression value over a control.

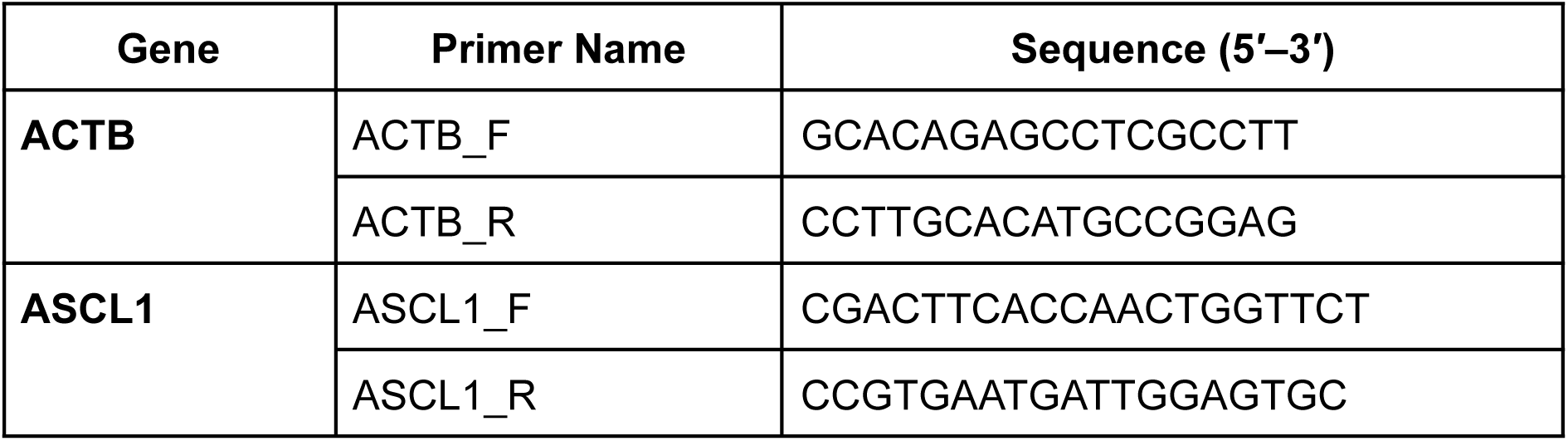
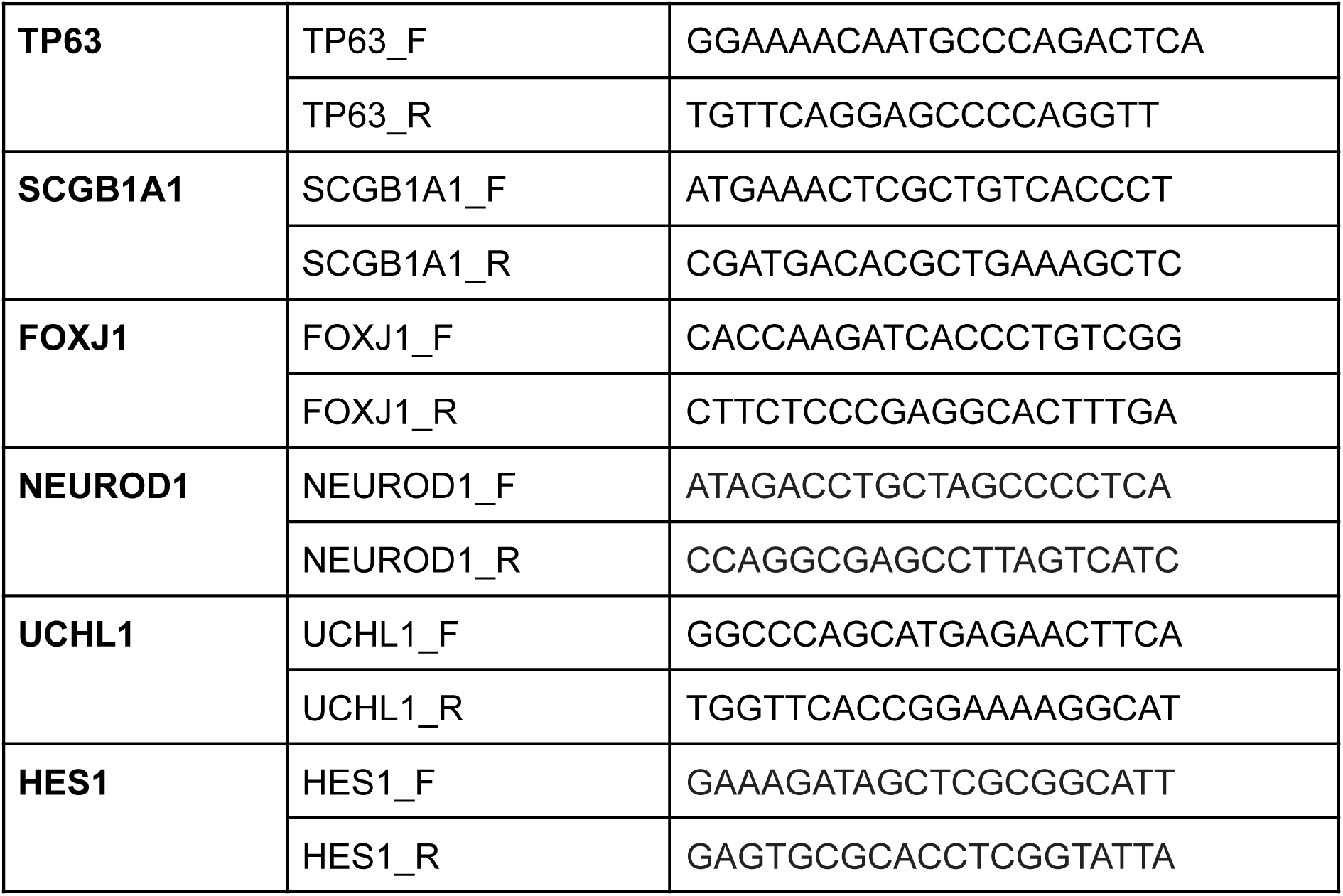

### scRNA-sequencing

*Lab work*. We performed single cell RNA sequencing (scRNA-seq) across 3 fAO lines in 2 different sequencing runs. First, 3 fAO lines were sequenced after confluent growth in NE expansion medium (15, 21, and 23 days, Table S1) and 2 of these lines were sequenced after confluent growth in standard media (8 and 21 days in culture, Table S1). In a subsequent set of runs, 3 fAO lines were expanded in NE expansion media for 14 to 22 days, at which point the media was switched to NE differentiation media and cells were collected after 1, 3, and 10 days of differentiation.

To dissociate organoids to single cell suspensions, BME drops were dissolved in 1–2 mL of TrypLE Express (Thermo Fisher Scientific, Cat# 12605010) and transferred to a 15 mL conical tube. Samples were incubated in a 37 °C water bath for 2-5 min, followed by mechanical shearing through a serological pipette fitted with a P10 tip to increase shear force. Cells were passed through a 35µm filter then spun down and resuspended in PBSA 0.1% for FACS sorting in 384-well plates (Single Cell Discoveries) for a SORT-seq2 workflow to generate barcoded single-cell libraries for transcriptome profiling. Library preparation and sequencing was performed by Single Cell Discoveries. In total 21 384-well plates were sequenced.

*Sequencing read alignments and quality control*. WGS raw sequencing reads generated from the SORT-seq2 workflow were processed using an in-house single-cell RNA-seq pipeline based on open-source tools. Initial quality assessment of raw FASTQ files was performed using FastQC (v0.12.1) to evaluate base quality scores, GC content, and adapter contamination. Adapter sequences and low-quality bases were trimmed using Skewer (v0.2.2) with default parameters, ensuring removal of short fragments prior to mapping. Filtered reads were aligned to the GRCh38 human reference genome with the gencode v44 comprehensive gene annotation as reference using STAR (v2.7.11a) in solo mode for single-cell data. The reference genome index was generated with STAR using the primary assembly FASTA and gencode annotation files. For barcode processing, cell barcode and unique molecular identifier (UMI) positions were specified according to the SORT-seq2 design, with error correction and multi-mapping handled through STAR’s internal algorithms. Alignments were output as coordinate-sorted BAM files with unmapped reads retained for downstream quality control. UMI collapsing, barcode demultiplexing, and gene count quantification were performed within STAR’s solo module to produce digital gene expression matrices, quantifying both spliced and unspliced reads using the Velocyto option, producing separate counts for spliced, unspliced, and ambiguous transcripts. This output was subsequently used for RNA velocity analyses to infer transcriptional dynamics.

*Single-cell data analysis.* All scRNA-Seq analysis downstream of gene expression matrices quantification was done using Scanpy^61^. For published datasets with existing cell-type annotations, we retained the cells included in the original analyses. For the unpublished fAOst dataset, quality control filtering removed potential multiplets (total counts >99.5th percentile), empty droplets (<1,000 counts), low-quality cells (<500 detected genes), and cells with high mitochondrial content (>25%). De-noised data matrix read counts per gene were log-normalized prior to analysis. For each cell, total counts across all genes were scaled to a common target sum (10,000 counts per cell), ensuring that all cells were comparable in overall expression levels, and Normalized counts were then log-transformed. After log-normalization, highly variable genes were identified and extracted. The normalized expression levels then underwent linear regression to remove effects of total reads per cell, mitochondrial genes, and cell cycle genes. Dimensionality reduction was performed using principal component analysis (PCA). Batch effects were corrected using the Harmony algorithm^62^, with sequencing sample ID specified as the batch variable. When integrating our SORT-seq dataset with the previously published 10x Chromium dataset, sequencing technology was additionally included as a covariate during PCA correction. Two-dimensional visualization was generated using a force-directed layout (FDL) algorithm applied to a 30-nearest-neighbor graph constructed from the top 25 principal components. Cell types were automatically annotated using CellTypist with a custom reference model trained on integrated single-cell atlases of human fetal and adult lung epithelium, followed by manual curation based on canonical marker genes. Clusters of cells within the fAOs and PNEC-extracted datasets were identified using the Leiden algorithm in Scanpy, with resolutions of 1.0 and 0.3, respectively. CellRank^38^ with RNA Velocity^60^ were used to infer directionality of cell state transition within PNEC population in an unbiased manner. We utilized default cell rank functions from Theis Lab (GitHub) to impute initial/terminal states, fate probabilities and TFs lineage drivers.

*Consensus scHPF.* Bayesian matrix factorization method that extends hierarchical Poisson factorization (HPF) to single-cell RNA-seq data, was trained on extracted basal cells using the consensus scHPF wrapper provided by the Sims Lab (GitHub). We included 20,920 feature genes, tested k=2–8 gene modules, ran t=20 trials, selected the top n=10 models per cluster, set a minimum of m=5 models per cluster, and allowed a maximum of j=10 models to run in parallel. Downstream analyses utilized cell scores for violin plots, correlation with single-cell–resolved gene signatures, and visualization via heatmaps and clustermaps annotated by cell type and media condition.

